# A binary arginine methylation switch on histone H3 Arginine 2 regulates its interaction with WDR5

**DOI:** 10.1101/2020.01.13.904581

**Authors:** Benjamin M. Lorton, Rajesh K. Harijan, Emmanuel S. Burgos, Jeffery B. Bonanno, Steven C. Almo, David Shechter

## Abstract

Histone H3 arginine 2 (H3R2) is post-translationally modified in three different states by “writers” of the protein arginine methyltransferase (PRMT) family. H3R2 methylarginine isoforms include PRMT5-catalyzed mono- and symmetric di-methylation (me1, me2s), and PRMT6-catalyzed me1 and asymmetric dimethylation (me2a). WD-40 repeat-containing protein 5 (WDR5) is an epigenetic “reader” protein that interacts with H3R2 and is a subunit of numerous chromatin-modifying complexes, such as the <u>M</u>ixed <u>L</u>ineage <u>L</u>eukemia (MLL) H3 lysine 4 methyltransferase complex. Previous studies suggested that MLL recruitment to chromatin was specified by the high-affinity interaction between WDR5 and H3R2me2s. However, our prior biological data prompted the hypothesis that WDR5 may also interact with H3R2me1 to recruit MLL activity. Here, using highly accurate quantitative binding analysis combined with high-resolution crystal structures of WDR5 in complex with unmodified (me0) and me1/me2s L-Arginine amino acids and in complex with H3R2me1 peptide, we provide a rigorous biochemical study of this important biological interaction. Despite modest structural differences at the binding interface, our study supports an interaction model regulated by a binary arginine methylation switch: H3R2me2a prevents interaction with WDR5, whereas H3R2me0/me1/me2s are equally permissive.

---

Arginine methylation (Rme) is a prevalent histone post-translational modification (PTM) that influences the ability of epigenetic “reader” proteins to engage with chromatin ^1^. By members of the protein arginine methyltransferase (PRMT) family, arginine can be monomethylated (Rme1, also MMA), asymmetrically dimethylated (Rme2a, also aDMA), or symmetrically dimethylated (Rme2s, also sDMA) ^2^. All PRMTs can generate Rme1; Type I PRMTs (isozymes 1-4,6,8) further catalyze Rme2a; Type II PRMTs (isozymes 5 and 9) further catalyze Rme2s; the sole Type III PRMT7 catalyzes only Rme1 ^3^. While each methylation event does not influence the arginine sidechain positive charge, it does reduce hydrogen bond (H-bond) capacity and alters the electronic distribution of the guanidino group by withdrawing electrons through hyperconjugation, thereby increasing hydrophobicity (Figure 1a) ^4^.

**Figure 1.**
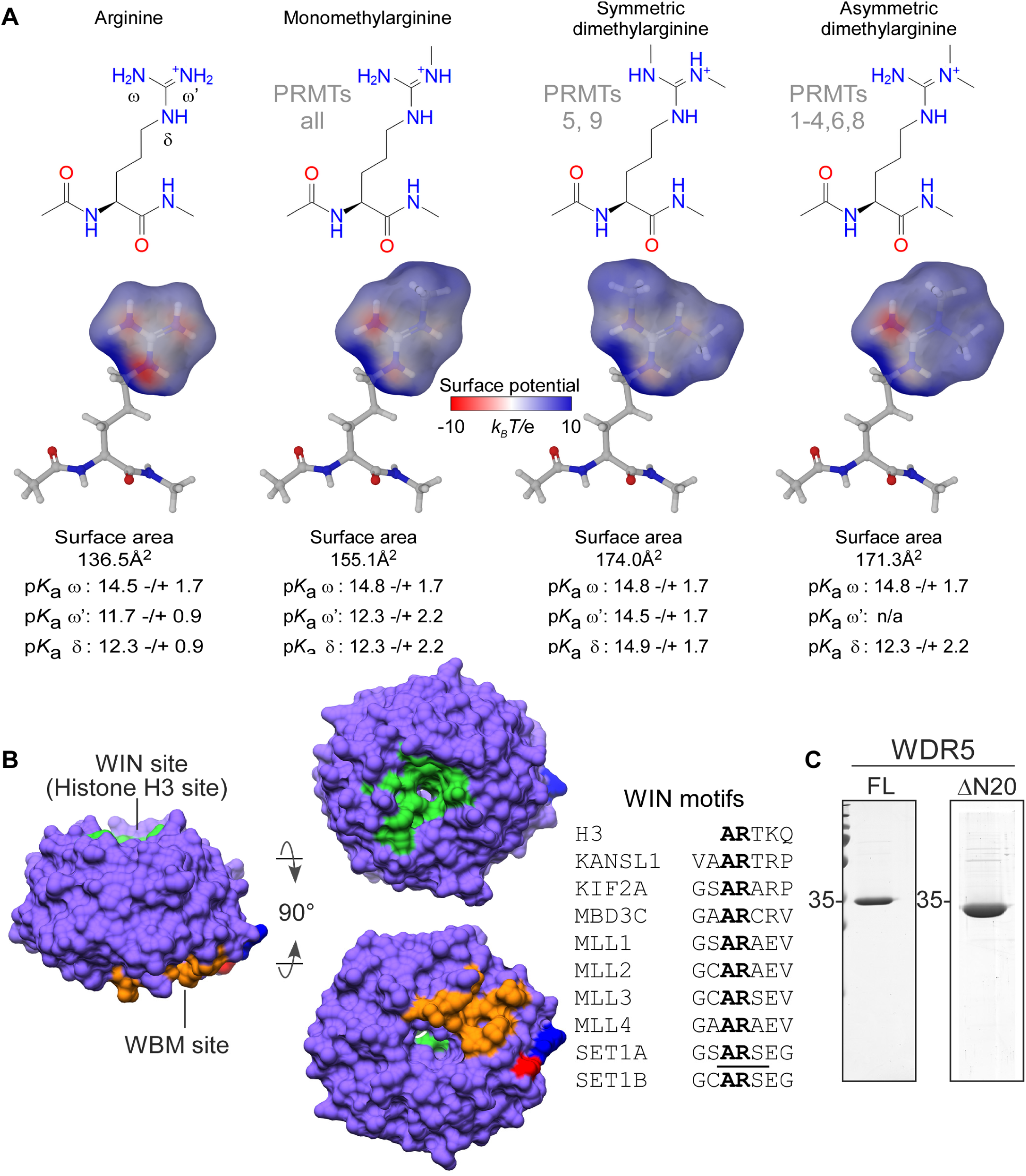
Overview of arginine methylation and WDR5. a. Guanidino group methylarginine isoforms which occur in vertebrates are depicted and annotated with their respective PRMT writer enzymes. Modeled surface areas and predicted guanidino group p*K*a values are shown (Maestro Schrödinger software) b. Surface representation of WDR5, depicting the WIN site (green), WBM site (orange), and the amino (blue) and carboxy (red) termini; characterized WIN motifs (underlined) are aligned c. Purified recombinant WDR5 proteins used in this study: full-length (FL) for all binding assays;1-20aa amino terminal truncation (ΔN20) for structural studies.

Although Rme has long been considered an irreversible PTM, emerging evidence suggests that a subset of lysine demethylase Jumonji-domain containing proteins catalyze arginine demethylation *in vitro* ^5^, suggesting that arginine methylation is reversible *in vivo*, where it is utilized to facilitate or disrupt protein-protein and protein-nucleic acid interactions that regulate diverse cellular processes ^6^. Histone arginine methylation is perhaps the most extensively studied methylarginine PTM for its role–in combination with other histone PTMs (e.g. lysine acetylation/methylation, serine/threonine phosphorylation)–in tuning chromatin state ^6^.

WD-40 repeat-containing protein 5 (WDR5) is a ubiquitously expressed and promiscuous adaptor protein that interacts with histone H3; it is also present in numerous chromatin-modifying complexes, including histone acetylases, deacetylases, and methylases ^7^. WDR5 is a globular protein with a narrow channel extending through the center of its seven WD-repeat β- propeller fold. Binding partners interact with WDR5 via two distinct and non-overlapping sites on opposite faces of the protein: the <u>W</u>DR5-<u>in</u>teraction (WIN) site and the WDR5 binding motif (WBM) site. Proteins that have been structurally characterized to interact with the WIN site contain a WIN motif: a four amino acid long polypeptide with consensus sequence A-R-[A/C/S/T]-[E/K/R] (Figure 1b, adapted from ^7^). The first four residues of histone H3 (A-R-T-K) comprise a WIN motif, and it is the only one known to be post-translationally modified with arginine methylation (PRMT6: me1, me2a ^8^; PRMT5: me1, me2s ^9^). Intriguingly, the sequence surrounding H3R8 (A-R-K-S) closely resembles and is hypothesized to constitute a second WIN motif on the H3 tail ^10–12^ that is also modified with arginine methylation (PRMT2: me1, me2a ^13^; PRMT5: me1, me2s ^14^).

Previously published structural studies revealed that WDR5 binds both unmodified H3R2 (H3R2me0) ^11, 15–17^ and H3R2me2s ^18^ peptides. It was previously reported that WDR5 is a high-affinity reader of H3R2me2s (*K*_d_∼100nM) ^18^ and, conversely, that H3R2me2a does not interact with WDR5 ^18–20^. This led to the hypothesis that WDR5 maintains open chromatin by interacting with H3R2me2s and recruits MLL methyltransferase activity to install H3 lysine 4 trimethylation (H3K4me3), potentiating transcription ^18^. As both H3R2me0 and H3R2me2s interact with WDR5, it follows that H3R2me1 can also be accommodated. However, in our search of the literature, we found only one study characterizing interaction with H3R2me1 that reported very weak affinity for WDR5 (*K*_d_ >500μM), which essentially excludes any biologically relevant interaction ^16^.

Rme1 has long been regarded as a non-functional transient species in the ‘obligatory’ pathways to either Rme2a or Rme2s ^21, 22^. However, numerous proteomic analyses demonstrate that Rme1 is an abundant PTM, suggesting it is functionally significant in the cell ^23–26^. Furthermore, we previously reported that TGF-ß stimulation of lung cancer cells resulted in a PRMT5-dependent increase in H3R2me1 at promoters and within gene bodies of monitored genes. PRMT5 knockdown or inhibition with GSK591 (a potent and specific PRMT5 inhibitor ^27^) resulted in depletion of H3R2me1 with concomitant loss of H3K4me3 and decreased expression of monitored genes. WDR5 knockdown or inhibition with OICR-9429 did not affect H3R2me1 levels but did significantly reduce H3K4me3 levels and reduced expression of assayed genes. These results led us to hypothesize that WDR5 can in fact read H3R2me1 to recruit MLL H3K4 methyltransferase activity to target genes and activate transcription ^9^.

To fully elucidate the biochemical details of this important biological relationship, the conflicting results in both biological and quantitative binding studies suggest that a more rigorous assessment of the WDR5/H3 interaction is warranted. Here, we present a complete interrogation of the H3/WDR5 interaction. Using extremely pure H3 peptides and highly accurate and precise peptide concentrations, as measured by 1D ^1^H <u>n</u>uclear <u>m</u>agnetic <u>r</u>esonance (NMR), we performed isothermal titration <u>c</u>alorimetry (ITC) to accurately measure the thermodynamic binding properties of the H3/WDR5 interaction. We show that full-length WDR5 (Figure 1c) specifically interacts with H3R2 and does not interact with H3R8. Furthermore, we provide unambiguous evidence that WDR5 indeed interacts with H3R2me1 and that it has comparable affinity for H3R2me0, H3R2me1, and H3R2me2s; additionally, we confirm no WDR5 interaction with H3R2me2a peptides. To assess structural differences at the WIN site, we solved high-resolution crystal structures of ΔN20-WDR5 (Figure 1c) in complex with H3R2me1 peptide and in complex with methylated isoforms of L-Arginine amino acid. We propose that the WDR5/H3R2 interaction is regulated by a binary methylation switch mechanism, such that H3R2me0/me1/me2s are equally permissive for, whereas H3R2me2a is excluded from, interaction with WDR5.

## Materials and Methods

### Expression and Purification of WDR5

Plasmids encoding human full-length WDR5 (2GNQ, Addgene) or a 20aa N-terminal truncation (ΔN20, prerequisite for crystallization; cloned into pRUTH5 expression vector)—both fused to an N-terminal 6x-Histidine affinity tag and Tobacco Etch Virus (tev) protease site— were transformed into *E. coli* BL21(DE3)pLysS. Cultures were grown in LB medium containing appropriate antibiotics at 37°C to OD600 ∼0.7 then induced with 1mM IPTG and incubated for 4 hours at 37°C. Per 1L worth of pelleted cells, lysis was performed by thorough resuspension in 35mL lysis/wash (LW) buffer (50mM Tris-HCl pH 8.0, 1M NaCl, 5mM β-mercaptoethanol, 5mM imidazole, 1mM PMSF). BugBuster (Millipore) was added to 1X and incubated at 25°C for 15min. This was followed by a sonication gradient: 2x30sec pulses at 35% and 40% amplitude and 2x10sec pulses at 45% and 50% amplitude with at least 30sec rest periods between pulses. Insoluble material was pelleted by centrifugation at 14,000rpm for 45min at 4°C. The soluble fraction was incubated with 1.25mL Ni-NTA affinity resin (Thermo) for 1hr with at 4°C. Ni-NTA gravity-flow columns were packed in 20mL disposable chromatography columns (BioRad) and washed with 50mL of buffer LW, 25mL LW+15mM imidazole, and then eluted with 5mL LW+360mM imidazole. Elution fractions containing 6xHis(tev)WDR5 were pooled, and 6x-Histidine-Tobacco Etch Virus protease catalytic domain was added at a 1:50 mass ratio (TEV:WDR5). TEV-cleavage was performed simultaneously with dialysis against 1L TD-buffer (50mM Tris-HCl pH 8.0, 200mM NaCl, 5mM ß-mercaptoethanol) for 16 hours at 4°C. TEV protease and the 6xHis affinity tag were then removed by subtractive Ni-NTA chromatography. To eliminate remaining contaminants, anion exchange chromatography was utilized (MonoQ, GE). In TD-buffer at pH 8.0, contaminants interact with MonoQ resin, whereas WDR5 (pI=8.1) does not; WDR5 was thus collected in the “unbound” fraction and contaminants eluted with a one-step increase to TD-buffer containing 1M NaCl. WDR5 was then polished by size exclusion chromatography (Superdex 75 Increase, GE) in TD-buffer containing 1M NaCl. Fractions containing pure WDR5 were pooled and buffer exchanged into storage buffer (20mM Tris-Cl pH 8.0, 150mM NaCl, 1mM TCEP). Full length WDR5 was concentrated to 500μM; ΔN20-WDR5 was concentrated to 660μM for crystallization experiments. Aliquots were stored at -80°C.

### Molecular Modeling and pKa Prediction of Methylarginine Isoforms

Maestro (Shrödinger) was used to model electrostatic surface potential maps and predict p*K*a’s of methylarginine isoforms. First, a G-R-G peptide was modeled using the “Build” feature in the software. At methylated positions of the arginine side chain, guanidino protons were changed to methyl groups using the “Set element” feature. Guanidino group atoms were then selected and electronic maps were generated using the “Poisson-Boltzmann ESP” task. Electrostatic potentials were displayed from -10 to 10 *k_B_*T/e (*k_B_*, Boltzmann constant; T, temperature (Kelvin); e, electron charge). Surface transparencies were set to 30% front and 10% back from the “Surface display options” menu. p*K*a values were predicted using Epik by selecting the “Emprical p*K*a” task.

### Histone H3 Peptide Pulldown Assay

10μg of C-terminal biotinylated histone H3 1-21aa unmodified, H3R2me1, H3R2me2s, or H3R2me2a peptide (Anaspec) were immobilized on 20μL Streptavidin sepharose resin (Thermo) in 200μL pulldown buffer (50mM Tris-Cl pH 8.0, 150mM NaCl, 5mM β-mercaptoethanol). The peptide-bound resin was washed three times with 500μL pulldown buffer then incubated with 10μM WDR5 full-length protein in 200μL buffer for 1hr with end-over-end rotation at 4°C. The resin was washed 8 times with 500μL buffer and eluted with 20μL 2X Laemelli Buffer and heat denatured at 90°C for 5 minutes. 15μL of eluate was loaded on 15% SDS-PAGE and visualized by Coomassie stain.

### Purification and Quantitative Analysis of Peptide Reagents

Histone H3 1-12aa unmodified, H3R2K, H3R8K, H3R2KR8K and H3 1-7aa unmodified, H3R2me1, H3R2me2s, and H3R2me2a peptides were obtained from GenScript USA. Extra care was exercised to ensure the highest purity and most accurate H3 peptide concentrations were obtained for quantitative binding measurements with WDR5. To this end, the peptides were again purified by <u>h</u>igh <u>p</u>erformance liquid <u>c</u>hromatography (HPLC), using a 250 x 4.6mm 5μm Luna C18-(2) column (Phenomenex; #00G-4252-E0) and a gradient separation with water (0.1% TFA; Phase A) and 90% acetonitrile (0.1% TFA; Phase B). All H3 1-12aa peptides were separated using the same protocol: 0-10 min (1 mL/min, 100% A), 10-40 min (1 mL/min, linear ramp to 75% A), 40-44 min (linear ramp to 100% B and 2 mL/min) with hold until 49 min, 49-53 min (2 mL/min, linear ramp to 100% A) with hold until 60 min. The H3 1-7aa peptides were separated following different protocols. Protocol #1 for H3R2me1 and H3R2me2s: 0-10 min (1 mL/min, 100% A), 10-90 min (1 mL/min, linear ramp to 85% A), 90-94 min (linear ramp to 100% B and 2 mL/min) with hold until 99 min, 99-103 min (2 mL/min, linear ramp to 100% A) with hold until 110 min. Protocol #2 for H3R2me0 and H3R2me2a: 0-10 min (1 mL/min, 100% A), 10-40 min (1 mL/min, linear ramp to 95% A), 40-44 min (linear ramp to 100% B and 2 mL/min) with hold until 49 min, 49-53 min (2 mL/min, linear ramp to 100% A) with hold until 60 min. Combined fractions of purified peptides were lyophilized and concentrations were determined using ^1^H NMR using an adenosine internal standard (D_2_O, 600 MHz, δ ppm: anomeric proton, ∼6.2, 1H, d, J=6.2 Hz) ^28^.

### Isothermal Titration Calorimetry

ITC experiments were performed using a PEAQ-ITC calorimeter (Malvern Panalytical). WDR5 aliquots were pooled and dialyzed against storage buffer. Lyophilized histone peptides and WDR5 were diluted into the filtered dialysate. WDR5 (100μM) was titrated with histone H3 peptide (1.5mM), deploying 19 injections (1x0.4μL, 18x2μL) at 25°C with a reference power of 2.5μcal/sec. Heat integration, baseline correction, and curve fitting were performed using the software accompanying the PEAQ-ITC. Stoichiometry was fixed at N=1 (one-site binding model), allowing for ΔH, *K*_d_, and displacement to vary for initial fitting iterations; the final iterations were performed with all variables free for determination of best fit. The best fit data were then imported into Prism to plot Langmuir isotherms.

### Crystallization of apo WDR5 and Liganded complexes

The crystallization of apo WDR5 was performed using sitting drop vapor diffusion method at 22°C. WDR5 was screened around previously published crystallization conditions ^18^. The crystallization drops were set up in 96-well INTELLI plates (Art Robbins) using the Gryphon crystallization robot (Art Robbins). Each crystallization drop contained 0.5μL of [28mg/mL] WDR5 and 0.5µL of well solution (0.1M Bis-Tris, 56.4mM ammonium sulfate, 32.5% PEG-3350). The volume of the well solution was 70µL. Diffraction quality crystals were obtained within one week. All liganded WDR5 complexes (unmodified, me1, me2s, and histone H3R2me1 1-21aa peptide) were obtained using crystal soaking experiments. The apo crystals of WDR5 were incubated for 30min at 25°C in soaking/cryoprotection buffer (well solution plus 20% glycerol), containing 2mM of L-Arg species or H3R2me1 1-7aa peptide. The crystallization and crystal handling processes are summarized in Table S2.

### Data collection and processing

The diffraction data were collected at LRL-CAT beam line (Argonne National Laboratory, Argonne, IL) at 0.97931Å wavelength. All diffraction data were processed using iMOSFLM ^29^ and scaled with the AIMLESS program of the CCP4 suite ^30^. Data quality was analyzed using SFCHECK and XTRIAGE ^30, 31^. Matthews coefficients (Vm) calculation were used to estimate the number of monomer molecules present the unit cells. The data collection and processing statistics are summarized in Table 1.

**Table 1.**
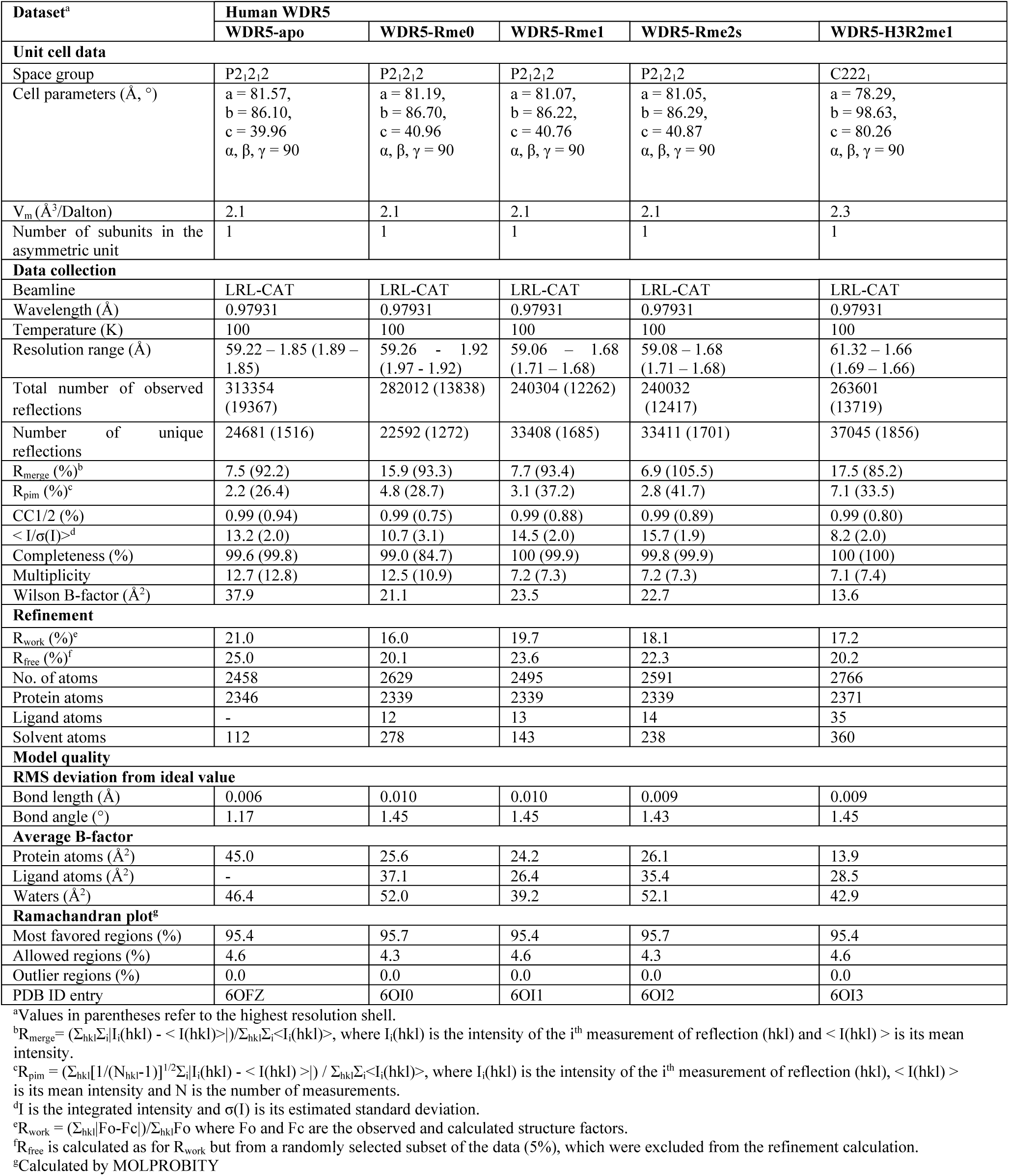
Data collection and refinement statistics of human WDR5.

### Structure determination

The crystal structures of WDR5 complexes were determined by molecular replacement using PHASER ^32^. Chain-A of wild-type WDR5 (PDB entry: 2H14) was used as the initial phasing template. The model obtained from PHASER was manually adjusted and completed using the graphics program COOT ^33^. Structure refinement was performed with the REFMAC5 program, using standard protocols for NCS refinement ^34^. Ligand molecules were omitted from the models in initial stages of building and refinement. After building all the water into the structures, L-Arg species molecules were fitted in their respective electron density features. The final refinement statistics of the structures are summarized in Table 1.

### Structure analysis

WDR5 subunit A from each structure was used as representative for all analyses and comparisons (A chain and B chain protein and ligand features were closely superimposable). Except for the images depicting modeled (methyl)arginine surfaces and ligand electron densities, which were generated using PyMOL (Schrödinger), all structural measurements and molecular graphic images were generated using the UCSF Chimera package from the Computer Graphics Laboratory, University of California, San Francisco (supported by NIH P41 RR-01081) ^35^. The geometry analyses of the final models were performed using MolProbity ^36^.

## Results

### Thermodynamic properties of WDR5 interaction with histone H3 peptides

Using an array of arginine-mutant and methylarginine-modified H3 peptides, we performed ITC experiments to completely interrogate the H3/WDR5 interaction. To provide the most accurate and precise affinity measurements, all commercially sourced peptides used in this study were: 1) repurified by high performance liquid chromatography (HPLC) and 2) concentrations were measured by NMR (Figure S1, S2, and Methods). All previously reported WDR5/H3R2 quantitative binding data were obtained using peptides containing H3R8 ^11, 15, 16, 18^. The amino acid sequence surrounding H3R8 (A-R-K-S) potentially constitutes a second WIN motif on the H3 tail that may have contributed to previously reported binding constants. Therefore, to interrogate WDR5/H3 interaction specificity, we performed replicate ITC experiments with H3 1-12aa (Figure 2a) wild-type and R-to-K mutant peptides: H3R2K, H3R8K, and H3R2KR8K.

**Figure 2.**
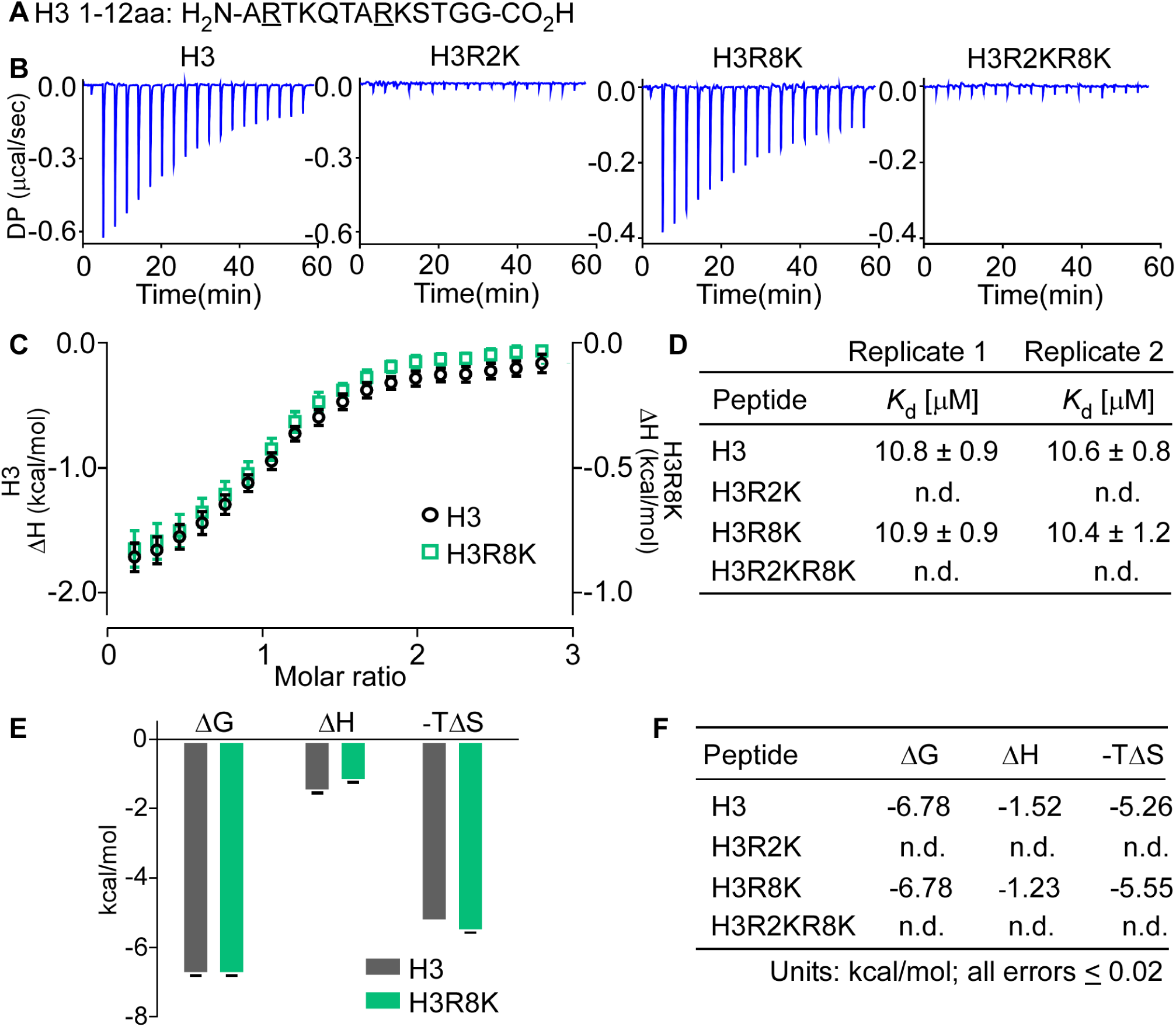
Replicate isothermal calorimetry assays showing WDR5 interacts with specifically with H3R2. a. Histone H3 1-7aa peptide sequence with methylarginine isoforms occurring at Arg2, underlined b. Raw power differentials (DP) of peptide titrations c. Integrated heats of binding; error bars represent the standard error between two replicate titrations d. Dissociation constants for replicate titrations determined by fitting heat integrations to a one-site binding model e. and f. Thermodynamic parameters associated with WDR5 interacting peptides (n.d., not detected) Peptides bound WDR5 with stoichiometries (N-values) between 0.9 and 1.1.

H3 wild-type and H3R8K peptides resulted in similar power differentials when injected into WDR5, whereas there was no indication of interaction with either the H3R2K or H3R2KR8K peptides (Figure 2b). Plotting enthalpy change (ΔH) versus H3 peptide:WDR5 (molar ratio) and fitting of a Langmuir isotherm (one-site binding model; Figure 2c) revealed that WDR5 has comparable equilibrium dissociation constants (*K*_d_; Figure 2d) for H3 (n=2; *K*_d_=10.8±0.9μM, *K*_d_=10.6±0.8μM) and H3R8K (n=2; *K*_d_=10.9±0.9μM, *K*_d_=10.4±1.2μM). Likewise, the thermodynamic parameters ΔG, ΔH, and -TΔS were similar between H3 and HR8K peptides Figure 2e,f). These results clearly demonstrate that WDR5 interacts specifically with H3R2 and does not interact with H3R8.

Next, to characterize the effect arginine methylation has on the H3R2/WDR5 interaction, we performed replicate ITC experiments with H3 1-7aa (Figure 3a) wild type or methylarginine-modified peptides: H3R2me1, H3R2me2s, H3R2me2a. Titration of H3R2me0, H3R2me1, and H3R2me2s 1-7aa peptides into WDR5 produced similar power differentials, whereas no heat was observed in titrations with H3R2me2a peptide (Figure 3b). These results are consistent with observations from our initial pulldown assays using H3 1-21aa peptides (Figure S3) and clearly demonstrate that WDR5 interacts with H3R2me0/me1/me2s but not with H3R2me2a. The data analysis using Langmuir isotherm (one-site binding model; Figure 3c) revealed that WDR5 has comparable equilibrium dissociation constants for H3R2me0 (n=2; *K*_d_ =11.0±0.5μM, *K*_d_ =11.8±0.5μM); H3R2me1 (n=2; *K*_d_ =9.0±0.8μM, *K*_d_ =8.1±1.4μM); and H3R2me2s (n=2; *K*_d_ =10.6±0.4μM, *K*_d_ =10.2±1.2μM), showing a modest increase in affinity for H3R2me1 (Figure 3d). Furthermore, the thermodynamic terms ΔG, ΔH, and -TΔS for each interaction are nearly identical (Figure 3e,f). With respect to H3R2me0/me1/me2s isoforms, our results establish that arginine methylation of H3R2 has a negligible effect on WDR5 interaction affinity and confirm no interaction with H3R2me2a. We were surprised that arginine methylation had such a small effect on WDR5 affinity and were thus compelled to investigate the structural differences between arginine- and methylarginine-liganded WDR5 complexes.

**Figure 3.**
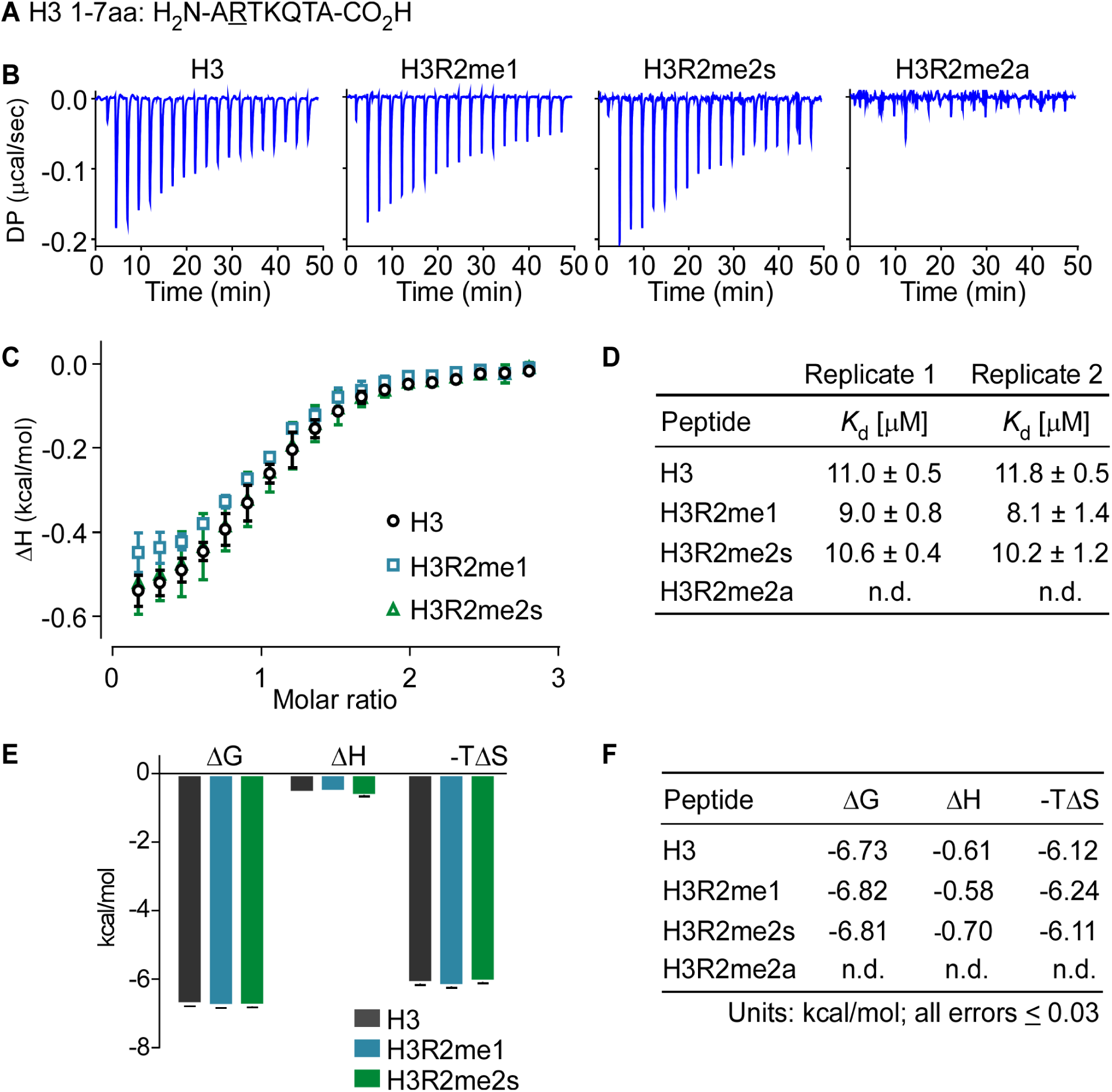
Replicate isothermal calorimetry assays showing WDR5 interacts with H3, H3R2me1, and H3R2me2s with similar heats and affinities but does not interact with H3R2me2a. a. Histone H3 1-7aa peptide sequence with methylarginine isoforms occurring at Arg2, underlined b. Raw power differentials (DP) of peptide titrations c. Integrated heats of binding; error bars represent the standard error between two replicate titrations d. Dissociation constants for replicate titrations determined by fitting integrated heats to a one-site binding model e and f. Thermodynamic parameters associated with WDR5 interacting peptides (n.d., not detected). Peptides that interacted with WDR5 bind with stoichiometries (N-values) between 0.9 and 1.

### Structure of WDR5 complexes

WDR5 is a ∼37kDa globular protein that adopts a 7-bladed ß-propeller fold (Figure 4a). At one end of the central channel is the WIN site, defined here as residues Ala47, Val48, Ser49, Ala65, Ser91, Glu107, Phe133, Cys134, Phe149, Phe173, Ser175, Tyr191, Tyr260, Cys261, Ile262, Phe263, Ile305, and Leu321 (Figure 1b). The solvent exposed outer surface of the WIN site is slightly acidic and becomes increasingly hydrophobic within the interior of the channel to accommodate the arginine sidechain (Figure S4). All structurally characterized WIN site ligands (Table S1) share a common binding mode: the WIN motif peptidyl arginine extends into the hydrophobic channel, with its guanidinium moiety stacking between aromatic residues Phe133 and Phe263 (Figure S5). As mentioned above, histone H3 is the only WIN site ligand known to be modified with methylarginine marks. Our binding studies establish that WDR5 has comparable affinity for H3R2me0/me1/me2s isoforms but has no interaction with H3R2me2a. To study any structural differences in binding the various methylarginine isoforms, high resolution crystal structures of unliganded (apo)WDR5 and WDR5 in complex with L-arginine (L-Arg) amino acid and its monomethyl (me1-L-Arg) and symmetric dimethyl (me2s-L-Arg) isoforms—completely devoid of additional peptide context—were determined. The structural analyses focused on the region of the WIN site interacting with the arginine side chain. The crystal structure of WDR5 bound to H3R2me1 peptide was also determined in the course of optimizing the system.

**Figure 4.**
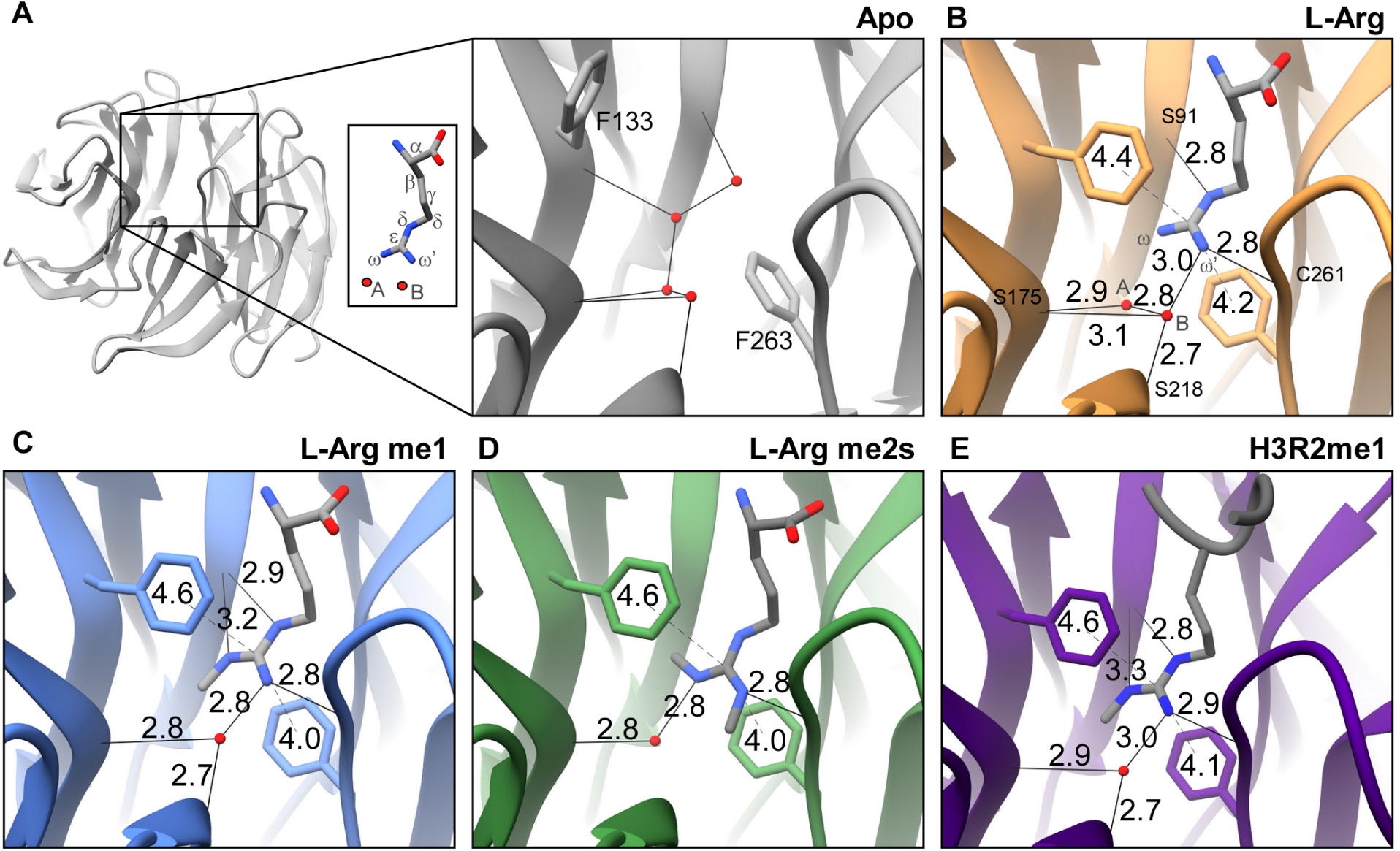
Crystal structures of the WDR5 WIN site detailing ligand guanidino group interactions with arginine and methylarginine isoforms. a. Overview of apo WDR5 with WIN site boxed and magnified. WDR5 complexes with b. L-Arg me0, c. L-Arg me1, d. L-Arg me2s, and e. H3R2me1 peptide ligands. Hydrogen bond (solid line) and p-stack (dashed line) distances are depicted. Water molecule (red sphere). Distances denoted in ångström (Å).

Crystallization of apo-WDR5 and conditions for obtaining WDR5 in complex with L-Arg, me1-L-Arg, and me2s-L-Arg amino acid or with H3R2me1 peptide are summarized in Table 2. To determine the crystal structures of the complexes, molecular replacement using PHASER was performed. WDR5 apo and L-Arg isoform complexed structures yielded data consistent with the P2_1_2_1_2 space group, whereas the WDR5/H3R2me1 peptide structure was determined in space group C222_1_—all structures contained one monomer in the asymmetric unit. The resolution of the structures varied between 1.66Å to 1.92Å. Except for a few amino acid residues at the amino and carboxy termini, electron density was well ordered over the entire WDR5 backbone structure. The sidechain electron density of several surface residues was also not resolved. All ordered amino acid residues were in the most favored or in additionally allowed regions of the Ramachandran plot (Table 1). The electron density corresponding to the L-Arg, me1-L-Arg, and me2s-L-Arg were well resolved, whereas the C-terminal residues of the H3 peptide were not observed in the structure (Figure S6).

**Table 2.**
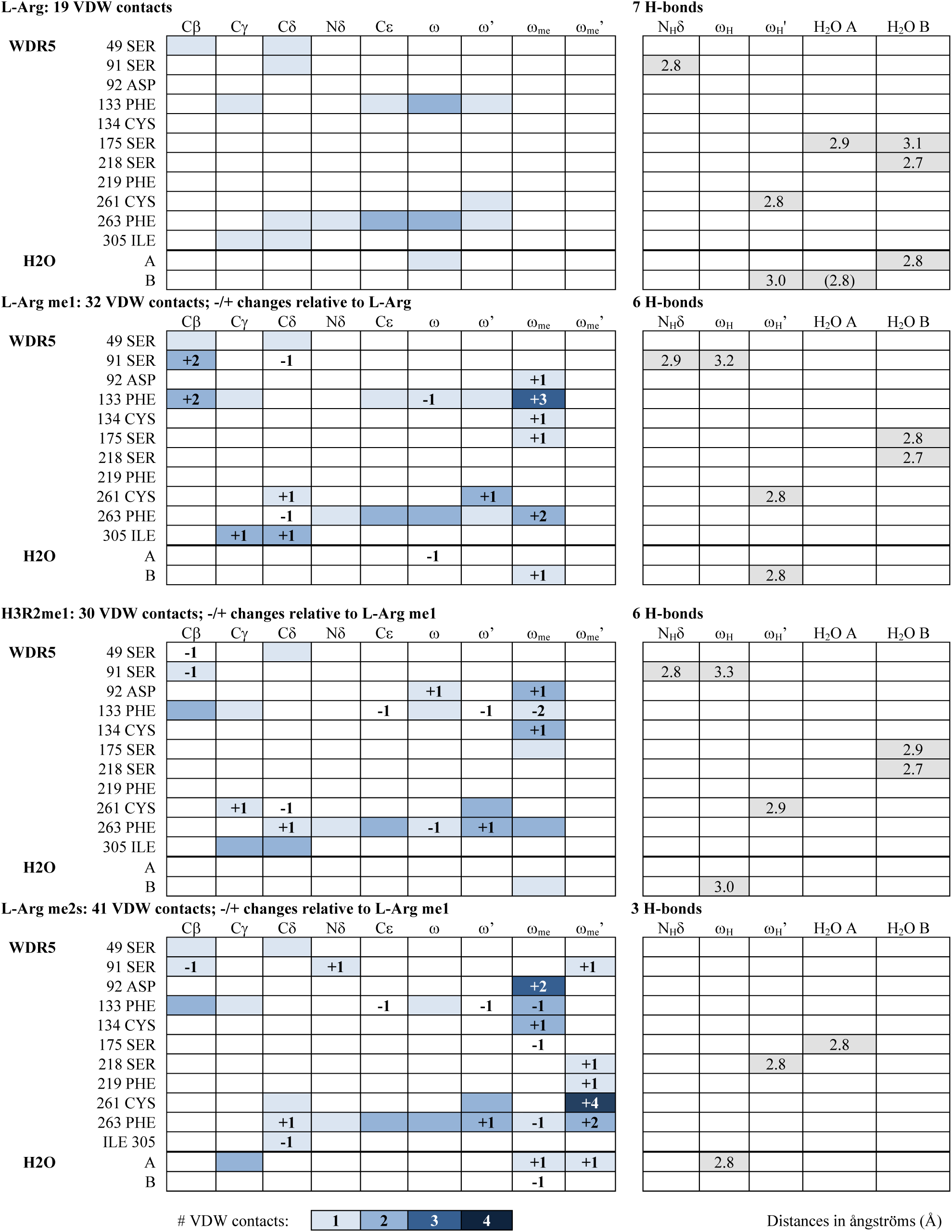
VDW contacts and H-bonds of liganded WDR5 complexes

### Binding of WDR5 with L-Arginine and methylarginine isoforms

Analysis of the WIN site of apo- and liganded-WDR5 structures provide details of the guanidino-binding interface. Hydrogen bonding, Van der Waals (VDW) contacts, and π-π stacking interactions were assessed in the structures (Figure 4, Table 2).

In the apo WDR5 structure, Phe133 and Phe263 aromatic sidechains are not in a stacked orientation, and the WIN site is occupied by four water molecules (Figure 4a). When bound to L-Arg ligand, Phe133 and Phe263 engage the guanidino group in π-π stacking interactions, and two waters remain in the WIN site (Figure 4b).These water molecules will be referred to as water A and water B, and the guanidino amino group proximal to water A will be referred to as ω and that proximal to water B as ω’ (Figure 4a, inset). A total of seven hydrogen bonds are formed between the arginine sidechain, structural water molecules, and WIN site residues in the L-Arg liganded structure (Figure 4b, Table 2). Water A forms one H-bond with the Ser175 backbone oxygen (O_B_) and one with water B. Water B is also hydrogen bonded to Ser175 O_B_ and to Ser218 O_B_. These four H-bonds are arranged in the same configuration in the apo structure, suggesting their structural significance (Figure 4a). The L-Arg guanidino group engages in three more H-bonds in the WIN site: one between L-Arg ω_H_’ and water B, another between L-Arg ω_H_’ and Cys261 O_B_, and a third between L-Arg N_H_δ and Ser91 O_B_.

We next inspected other contacts within VDW-interaction distances (Figure S7); WDR5 residues are listed with corresponding L-Arg atom interactions in Table 2. With respect to L-Arg, 19 VDW contacts are observed in the WIN site: 18 with WDR5 residues and one with water A (Figure S7a).

The aromatic rings of Phe133 and Phe263 engage in π-π stacking interactions with the delocalized electrons of guanidino group atoms. To assess whether arginine methylation affects this interaction, we measured the distance from the center of each phenylalanine aromatic ring to the guanidinium group central carbon (C_ε_) for each complex (Figure 4). With respect to L-Arg, the distance from Phe133−L-Arg was 4.4Å and Phe263−L-Arg was 4.2Å, giving a total π-π stacking distance of 8.6Å (Figure 4b).

When me1-L-Arg occupies the WIN site, the bulk of the guanidino methyl group (ω_me_) results in water A being displaced from the binding site (Figure 4c), resulting in a net loss of one hydrogen bond compared to the L-Arg structure. Water B remains engaged in the same 3 H-bonds as described for L-Arg; however, the hydrogen bond between me1-L-Arg ω_H_’ and water B and that between water B and Ser175 O_B_ are slightly shorter, and thus stronger. The hydrogen bond interaction between Ser91 O_B_ and N_H_δ is slightly longer, and one additional H-bond is gained between Ser91 O_B_ and me1-L-Arg ω_H_. 31 VDW contacts are formed between me1-L-Arg and WIN site residues and one with water B—13 more than L-Arg (Table 2, Figure S7b). The overall π-π stacking distance is unchanged (Figure 4c); however, the distance between Phe133−me1-L-Arg (4.6Å) is slightly increased by 0.2Å and the distance between Phe263−me1-L-Arg (4.0Å) is decreased by 0.2Å, compared to L-Arg (4.4Å and 4.2Å, respectively).

When WDR5 binds me2s-L-Arg, the guanidino moiety is rotated 180° about the Nδ-C_ε_ bond, such that the ω_me_ group is arranged perpendicular relative to its position in the me1-L-Arg structure. This arrangement now allows water A to be retained in the WIN site, whereas the bulk of me2s-L-Arg ω_me_’ results in the displacement of water B (Figure 4d). Overall, this results in a net loss of 3 H-bonds compared to the me1-L-Arg structure. Water A forms a slightly shorter H-bond with Ser175 O_B_ and another with ω_H_. The two hydrogen bonding interactions between me1-L-Arg ω_H_ and Ser91 O_B_ and between me1-L-Arg N_H_δ Ser91 O_B_ are lost in the me2s-L-Arg structure. 39 VDW contacts are made between me2s-L-Arg and WDR5 and one with both water A and water B—nine more than compared to the me1-L-Arg structure (Table 2, Figure S7c). The total π-π stacking distance is unchanged at 8.6Å (Figure 4d).

### Binding of WDR5 with histone H3R2me1 peptide

The crystal structure of WDR5 in complex with H3R2me1 1-21aa peptide was determined at 1.66Å resolution. The four N-terminal residues of the peptide are clearly visible at the binding pocket, whereas the remaining carboxy terminal residues were not resolved and presumably disordered. The arginine sidechain of the H3R2me1 peptide interacts with WDR5 much like Rme1 with only modest differences (Figure 4e). Peptidyl H3R2me1 makes 29 VDW contacts with WDR5 and one with water B—two fewer compared to me1-L-Arg (Table 2, Figure S7d). The π-π stacking interaction distance totals 8.7Å; the distance between Phe133−H3R2me1 is unchanged, whereas the distance between Phe263−H3R2me1 is slightly increased by 0.1Å to 4.1Å (Figure 4e).

### Structural comparisons

The overall Cα-Cα RMSD between apo and liganded structures is 0.381Å (L-Arg), 0.359Å (L-Arg me1), 0.324Å (L-Arg me2s), and 0.326Å (H3R2me1 peptide). Additionally, the RMSD between common interacting residues (Ser49, Ser91, Asp92, Phe133, Cys134, Ser175, Ser218, Phe219, Cys261, and Phe263) of liganded structures are all less than 0.22Å,, indicating strong agreement with very little deviation between structures. Comparing B-factors of the WDR5 WIN site bound to the L-Arg isoforms shows that each complex is of similar rigidity with B-factors ranging between 15-20Å^2^ (Figure S8). The miniscule differences in RMSD values and B-factors are consistent with our binary methylarginine switch interaction model: WDR5 does not discriminate between unmodified, mono-, and symmetric dimethyl-arginine ligands (Figure 5).

**Figure 5.**
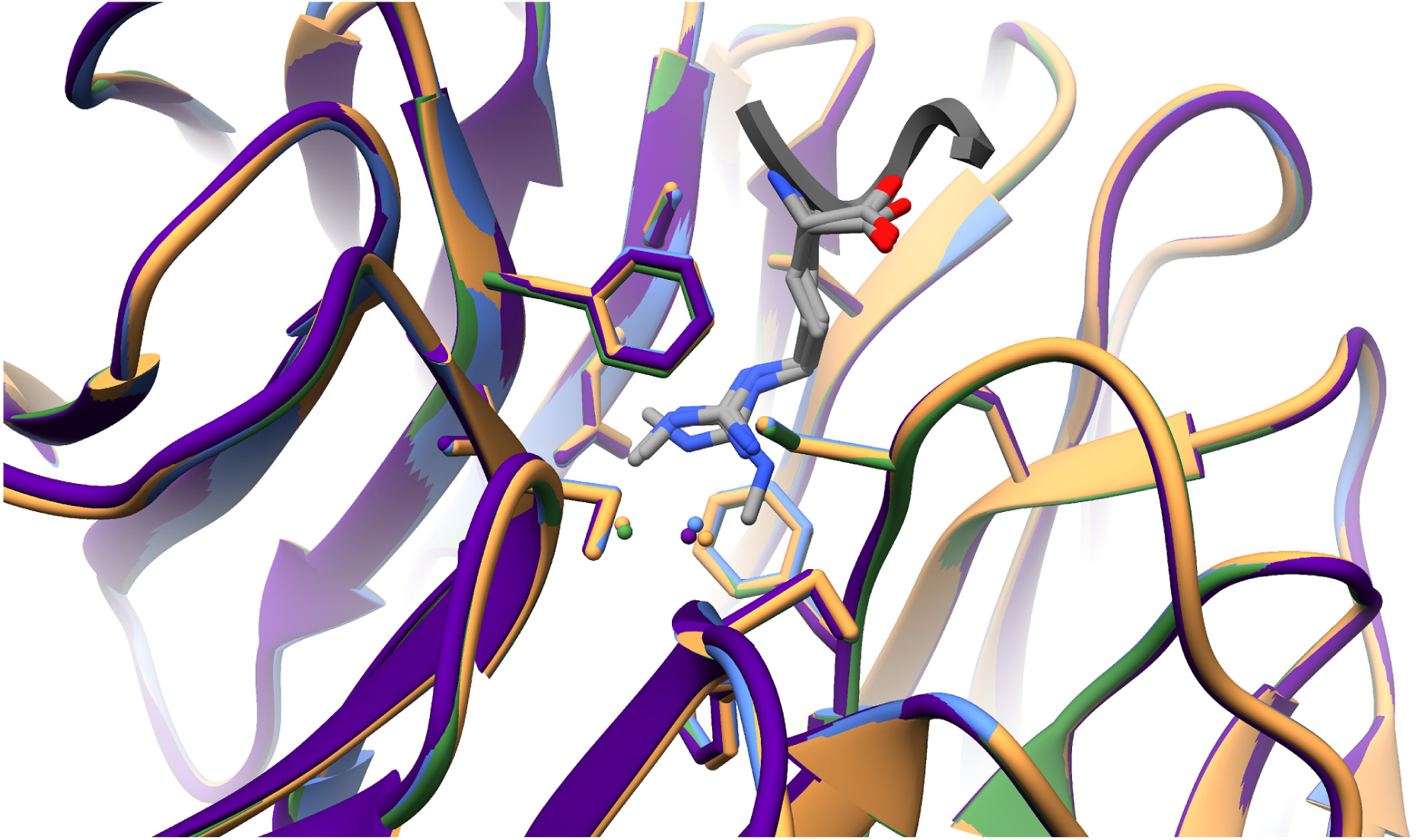
Superimposition of L-Arg(me) isoforms- and H3R2me1 peptide-liganded WDR5 complexes showing very modest differences between structures. WDR5 backbone and Arg-interacting residues are shown, with water molecules (spheres) colored accordingly. L-Arg (tan); L-Arg me1 (light blue); L-Arg me2s (green); H3R2me1 peptide (purple).

Consistent with our observations that Rme2a does not interact with WDR5, we did not observe electron density in the WIN site of WDR5 crystals after soaking with asymmetric dimethyl-L-arginine (Rme2a). Modelling Rme2a into the WDR5 WIN site revealed clashes that potentially prevent interaction (Figure S9). Our attempts to crystallize WDR5 bound to H3R8 using the H3R2me2a 1-21aa peptide were also unsuccessful.

## Discussion

For many years, the WDR5/H3 interaction has been of considerable interest for its role in regulating gene expression and development ^12^. However, previously published biological ^9^ and quantitative binding ^16^ data disagree with respect to which H3R2 methylarginine isoforms interact with WDR5. To answer this open question, we provide a thorough and rigorous biochemical assessment of the WDR5/H3 N-terminal tail interaction. We unequivocally demonstrate that WDR5 binding is: 1) specific for H3R2, showing no interaction with H3R8; and 2) is regulated by a binary methylarginine switch mechanism. In this binary switch, H3R2me0/me1/me2s all have near equal affinity for WDR5, whereas H3R2me2a is not compatible for interaction.

We utilized ITC with a variety of H3 peptides to interrogate the thermodynamics of interaction with WDR5. A major technical challenge to quantitatively probing histone tail interactions is the lack of aromatic residues, making measurement of peptide concentration by A280 impossible. Furthermore, although peptide concentrations can be measured by A205, this method relies either on a standard ^37^ or predicted ^38, 39^ molar extinction coefficient—both of which are estimated and introduce inaccuracies in final reported quantitative binding constant values. Additionally, we wanted to ensure that our peptides were of the utmost purity in our experiments. To address these concerns, we first repurified the peptides by HPLC over a much longer elution gradient than that normally used by peptide suppliers. We then measured peptide concentration using NMR, which is considerably more sensitive than A205 and allows for both highly accurate and precise concentration measurements.

WDR5 interacting proteins contain a WIN-motif—a four amino acid peptide with a consensus sequence of A-R-(A/C/S/T)-(E/K/R). The first two WIN motif residues are completely conserved with some plasticity at positions three and four. Histone H3 possesses a WIN motif at 1ARTK4 and is hypothesized to carry another at 7ARKS10 ^10–12^—both of which are modified by PRMT 5 and 6 and PRMT 2 and 5, respectively. Therefore, to delineate which H3 arginine residues interact with WDR5, H3 1-12aa peptides were synthesized and each arginine was systematically mutated to lysine followed by ITC measurements to quantify binding. Our results demonstrate that WDR5 only interacts with peptides containing H3R2; when H3R2 is mutated to lysine, no interaction is detected although H3R8 remains available for binding. These results define WIN motif requirements, i.e. position three requires a small R-group amino acid, and position four requires a charged sidechain. Furthermore, this offers a remarkable example of how a simple substitution of residues at WIN motif positions three and four imparts WDR5 binding specificity without post-translational modification.

WDR5 has been reported as interacting with H3R2me0 ^11, 15–17^ and H3R2me2s ^18^, with the latter described as a high affinity interaction *in vitro* ^18^. Both H3R2me1 ^16^ and H3R2me2a ^18–20^ had previously been reported not to interact with WDR5. However, more recent biological data from our group suggested that WDR5 does interact with H3R2me1 ^9^. Therefore, to address the discrepancies between biological and biochemical reports, we synthesized H3 1-7aa peptides containing the full array of H3R2 methylarginine isoforms and performed ITC to determine which isoforms interact with WDR5. Our results clearly show that WDR5 indeed interacts with H3R2me1 and, furthermore, that H3R2me0/me1/me2s all bind WDR5 with near equal affinity, whereas H3R2me2a is excluded. In line with findings that WDR5 indeed interacts with monomethylarginine, small molecule WIN site inhibitors have been designed that utilize a monomethyl-like appendage to engage WDR5 ^40, 41^. For all interacting H3 peptide ligands, the energy required is ∼90% entropic, indicating that binding is predominantly driven by a relatively large solvent gain upon rearrangement of the WDR5 WIN site and ligand arginine hydration shells (i.e. hydrophobic effect) with a small change in bond enthalpy. These results reveal that WDR5/H3R2 interactions are governed by a binary methylarginine switch, such that H3R2me0/me1/me2s are equally favorable for binding, whereas H3R2me2a is not.

As unmodified arginine and its methylated isoforms are structurally distinct, we attempted to assess the differences among the ligands when complexed with WDR5. To that end, we determined high-resolution crystal structures of WDR5 in complex with unmodified and methylated (me1, me2s) L-Arg amino acids and in complex with H3R2me1 peptide. Unmodified L-Arg interacts with WDR5 by utilizing two structural water molecules, seven H-bonds, and 19 VDW contacts. When me1-L-Arg binds WDR5, one water molecule is displaced which results in the net loss of one H-bond, but the bulk of the methyl group increases VDW contacts to 32. When me2s-L-Arg binds WDR5, the opposite water molecule is displaced which results in the loss of three more H-bonds; with two methyl groups present, VDW contacts are increased to 41. Arginine methylation had no effect on the π-π stacking interaction distance made with Phe133 and Phe263. Additionally, comparison of the RMSD values and B-factors of each liganded WDR5 complex shows that no major differences are brought about by arginine methylation. Thus, although arginine methylation decreases WIN site hydrogen bonding interactions, the loss of associated binding energy is compensated by an increase in VDW contacts. This compensation is consistent with the low change in enthalpy observed in our quantitative binding analysis and further supports that guanidino-group ligand interactions with WDR5 are primarily entropic: binding is driven by hydrophobic π-π interactions between Phe133, Phe263, and the ligand guanidino moiety and by desolvation of the arginine sidechain and WIN site pocket. Liberation of a water molecule from a protein cavity is predicted to contribute -0.46 to -2.67 kcal/mol of entropy (-TΔS) to binding interactions, with higher values associated with water molecules near charged residues ^42^. As the interior WIN site cavity is hydrophobic and the unbound arginine sidechain, although charged, is not buried, the entropy gained by liberation of each water molecule is likely toward the lower end of this scale. Our data is consistent with the displacement of 2-3 water molecules from the WIN site and another handful from the arginine sidechain upon WDR5/H3R2 interaction.

WDR5 interacts with transcription factors, such as MYC, and is present in numerous chromatin-modifying complexes, such as KANSL (lysine Acetyltransferase Non-Specific Lethal); NuRD (Nucleosome Remodeling and Deacetylase); and MLL ^7^. Moreover, WDR5 is expressed at levels considerably higher than other components of these complexes in vivo. It is enticing to speculate that WDR5 is stored on and maintains open chromatin at sites of unmodified, monomethyl, and symmetric dimethylarginine. As epigenetic complexes generally carry multiple reader subunits that stabilize interactions with chromatin, it raises the possibility that chromatin bound WDR5 may serve as a homing beacon for partially assembled complexes, thereby completing assembly and activating complexes as WDR5 disengages chromatin. Consistent with this hypothesis, two recent cryo-EM structures of MLL were determined in complex with the nucleosome core particle: one structure showed WDR5 engaged with the MLL WIN motif ^43^, whereas the WDR5 WIN site is unoccupied in the other ^44^.

In conclusion, this study clearly demonstrates that WDR5 interacts equally with unmodified, monomethyl-, or symmetric dimethyl-arginine but does not interact with asymmetric dimethylarginine. Moreover, this binary switch mechanism suggests that the purpose of installing either dimethylarginine isoform is to prevent installation of the other isoform, thereby creating or blocking docking sites for interacting proteins, which may serve as a global regulatory mechanism for WDR5-containing and/or other methylarginine-reading complexes.

---

**Supplementary Table 1.**
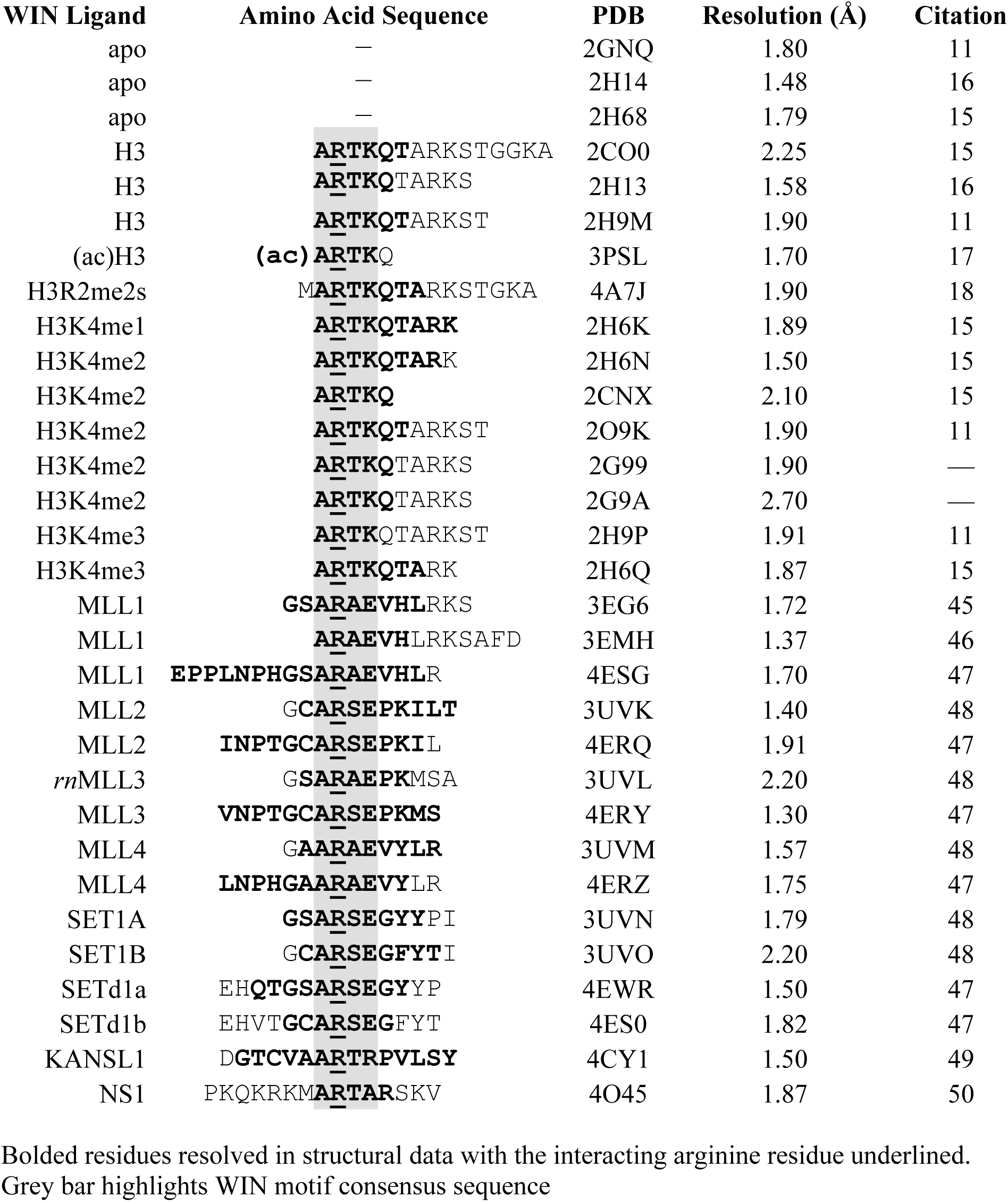
Alignment of WIN site ligands in solved WDR5 structures

**Supplementary Table 2.** WDR5 crystallization and crystal handling

| **WDR5 crystals** | **Crystallization conditions** | **Soaking/Cryoprotectant Solution** | **Space group** | **PDB ID** |
| --- | --- | --- | --- | --- |
| **WDR5-**<br>**apo** | 100 mM Bis-Tris pH5.9<br>32% PEG3350<br>54.6 mM Ammonium Sulfate | 100 mM Bis-Tris pH5.9<br>32.5% PEG3350<br>54.6 mM Ammonium Sulfate<br>20% Glycerol | P2<sub>1</sub>2<sub>1</sub>2 | 6OFZ |
| **WDR5-**<br>**Rme0** | 100 mM Bis-Tris pH5.9<br>32% PEG3350<br>54.6 mM Ammonium Sulfate | 100 mM Bis-Tris pH5.9<br>32.5% PEG3350<br>54.6 mM Ammonium Sulfate<br>20% Glycerol<br>2mM L-Arg | P2<sub>1</sub>2<sub>1</sub>2 | 6OI0 |
| **WDR5-**<br>**Rme1** | 100 mM Bis-Tris pH5.9<br>32% PEG3350<br>54.6 mM Ammonium Sulfate | 100 mM Bis-Tris pH5.9<br>32.5% PEG3350<br>54.6 mM Ammonium Sulfate<br>20% Glycerol<br>2mM L-Arg me1 | P2<sub>1</sub>2<sub>1</sub>2 | 6OI1 |
| **WDR5-**<br>**Rme2s** | 100 mM Bis-Tris pH5.9<br>32% PEG3350<br>54.6 mM Ammonium Sulfate | 100 mM Bis-Tris pH5.9<br>32.5% PEG3350<br>54.6 mM Ammonium Sulfate<br>20% Glycerol<br>2mM L-Arg me2s | P2<sub>1</sub>2<sub>1</sub>2 | 6OI2 |
| **WDR5-**<br>**H3R2me1** | 100 mM Bis-Tris pH5.9<br>32% PEG3350<br>54.6 mM Ammonium Sulfate | 100 mM Bis-Tris pH5.9<br>32.5% PEG3350<br>54.6 mM Ammonium Sulfate<br>20% Glycerol<br>2mM H3R2me1 1-21aa peptide | C222<sub>1</sub> | 6OI3 |

## ACCESSION NUMBERS

The structural data supporting this study have been deposited in the PBD. The accession numbers for each structure are: apo-WDR5 (6ofz); L-Arg/WDR5 (6oi0); me1-L-Arg/WDR5 (6oi1); me2s-L-Arg/WDR5 (6oi2); and H3R2me1/WDR5 (6oi3). The UniProtKB accession number for human WDR5 is P61964.

## ASSOCIATED CONTENT

The following files are available free of charge. Supplementary Figures 1-9 (PDF)

## AUTHOR INFORMATION

### Author Contributions

B.M.L. conceived the hypothesis and the experiments, performed all the studies, and wrote the manuscript; R.K.H. and J.B.B. determined crystal structures; E.S.B. assisted with Analytical Chemistry of peptides required for binding studies (HPLC purification and NMR quantification); S.C.A. supervised crystallography; D.S. supervised all studies and wrote the manuscript. All authors reviewed the final manuscript.

Conflict of interest: None declared.

## ACKNOWLEDGMENT

We would like to thank Maxim Maron for thoroughly reading and commenting on the final manuscript.

## ABBREVIATIONS

H3: histone H3
HPLC: high performance liquid chromatography
ITC: isothermal titration calorimetry
K4: lysine 4
*K*_d_,: equilibrium dissociation constant
KANSL: lysine acetyltransferase non-specific lethal
MLL: mixed lineage leukemia
NuRD: nucleosome remodeling and deacetylase
PRMT: protein arginine methyltransferase
R2: arginine 2
Rme1/MMA: monomethylarginine
Rme2s/sDMA: symmetric dimethylarginine
Rme2a/aDMA: asymmetric dimethylarginine
RMSD: root mean squared deviation
WBM: WDR5 binding motif
WDR5: WD-40 repeat-containing protein 5
WIN: WDR5 interacting.

---

**Figure S1.**
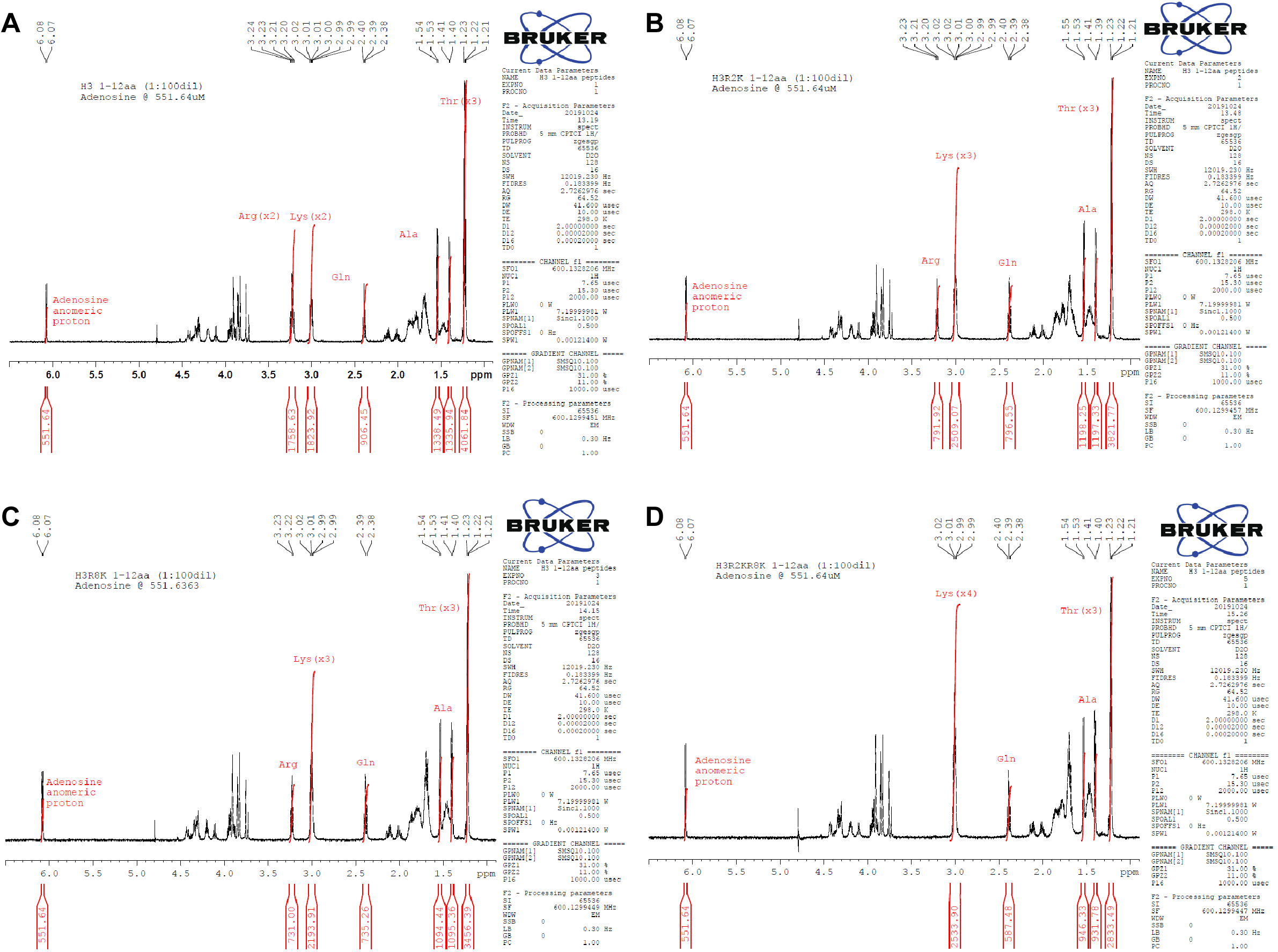
1D ^1^H NMR spectra of H3 1-12aa peptides showing peak integrations of residues used to measure peptide concentration. **a.** H3 unmodified **b.** H3R2K **c.** H3R8K. **d.** H3R2KR8K

**Figure S2.**
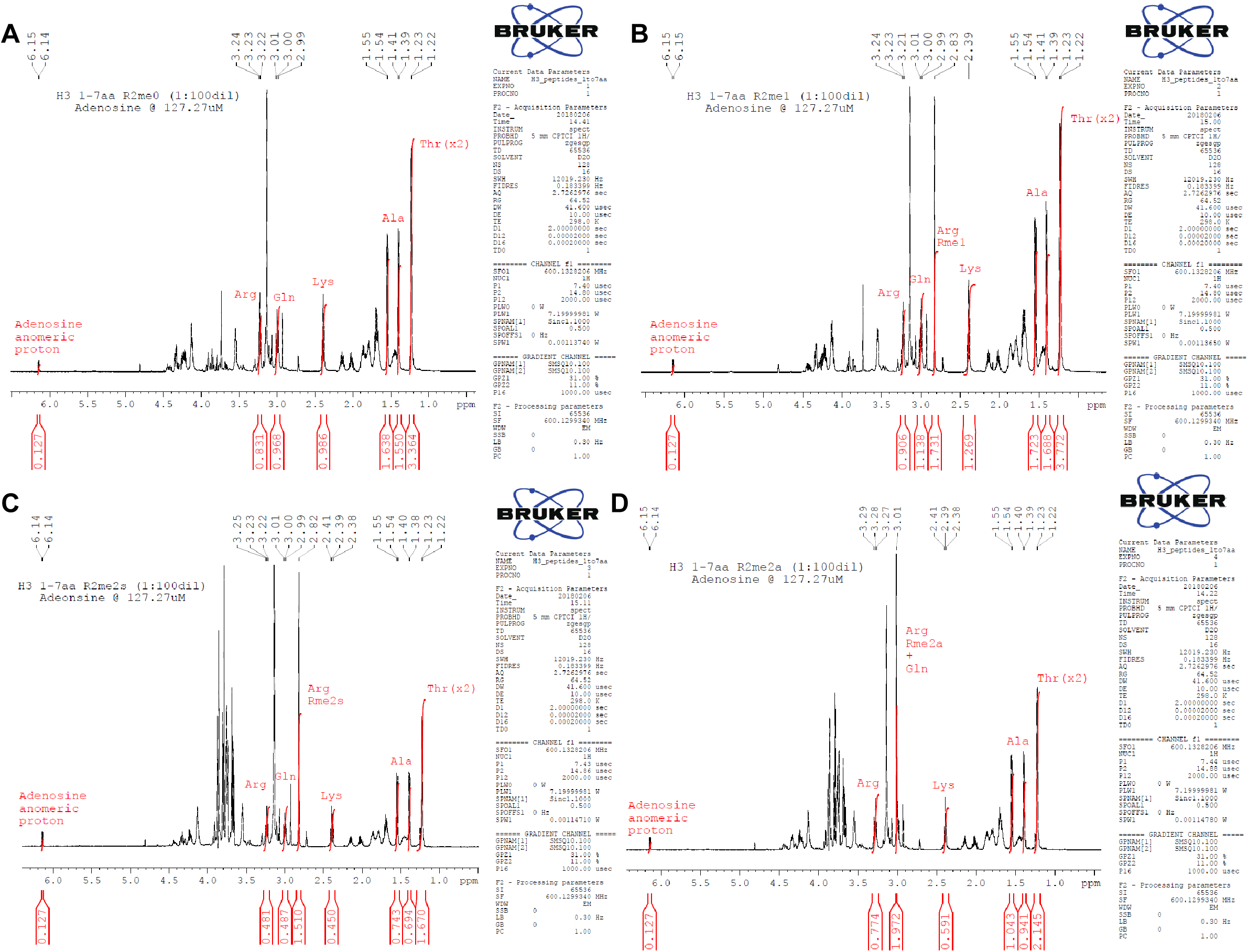
1D ^1^H NMR spectra of H3 1-7aa peptides showing peak integrations of residues used to measure peptide concentration. **a.** H3 unmodified **b.** H3R2me1 **c.** H3R2me2s. **d.** H3R2me2a

**Figure S3.**
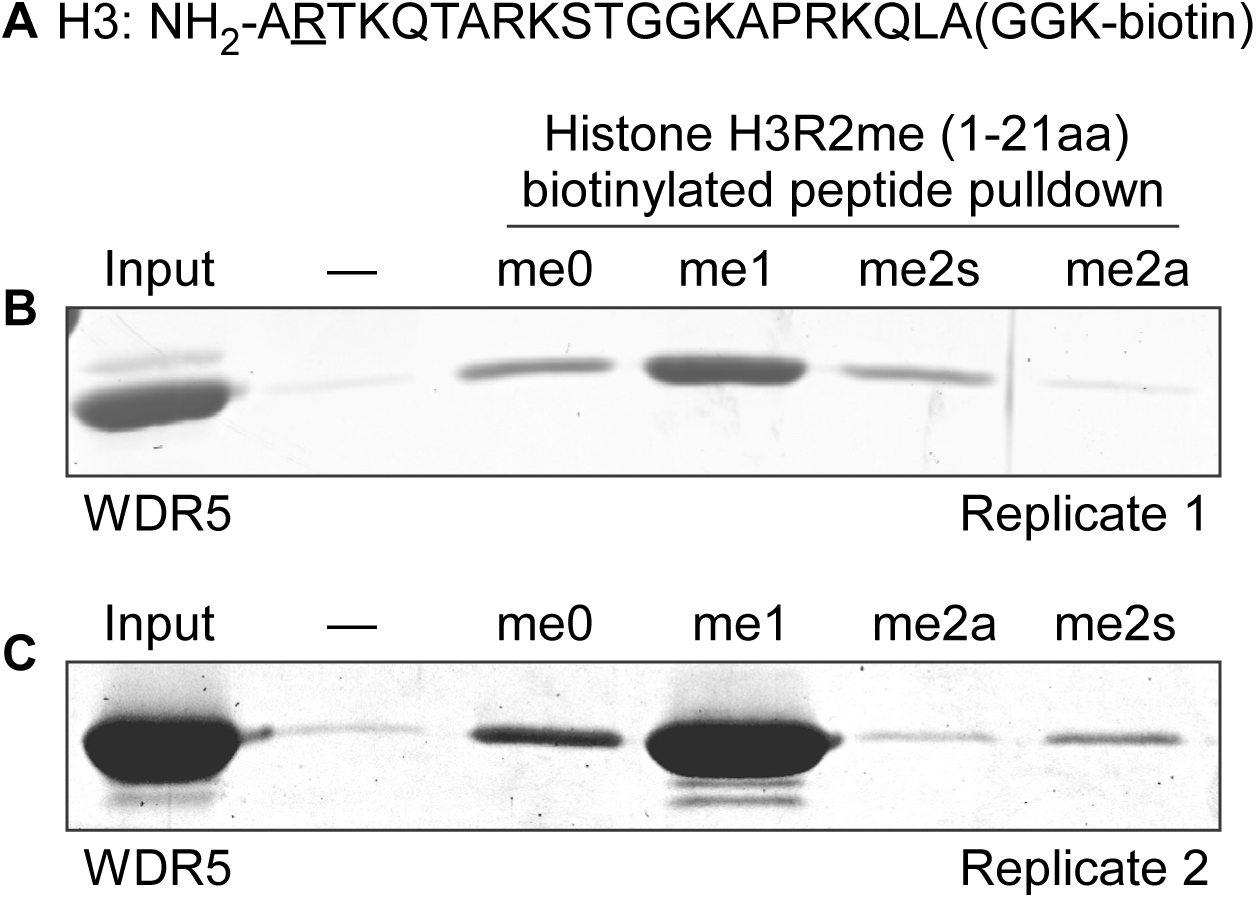
Replicate peptide pulldown assays showing WDR5 interacts with H3R2me0, me1, and me2s but not H3R2me2a. (-) negative control: no peptide, resin only. **a.** Histone H3 1-21aa peptide sequence with methylarginine isoforms ocurring at Arg2, underlined. **b.** and **c.** Replicate pulldowns 1 and 2

**Figure S4.**
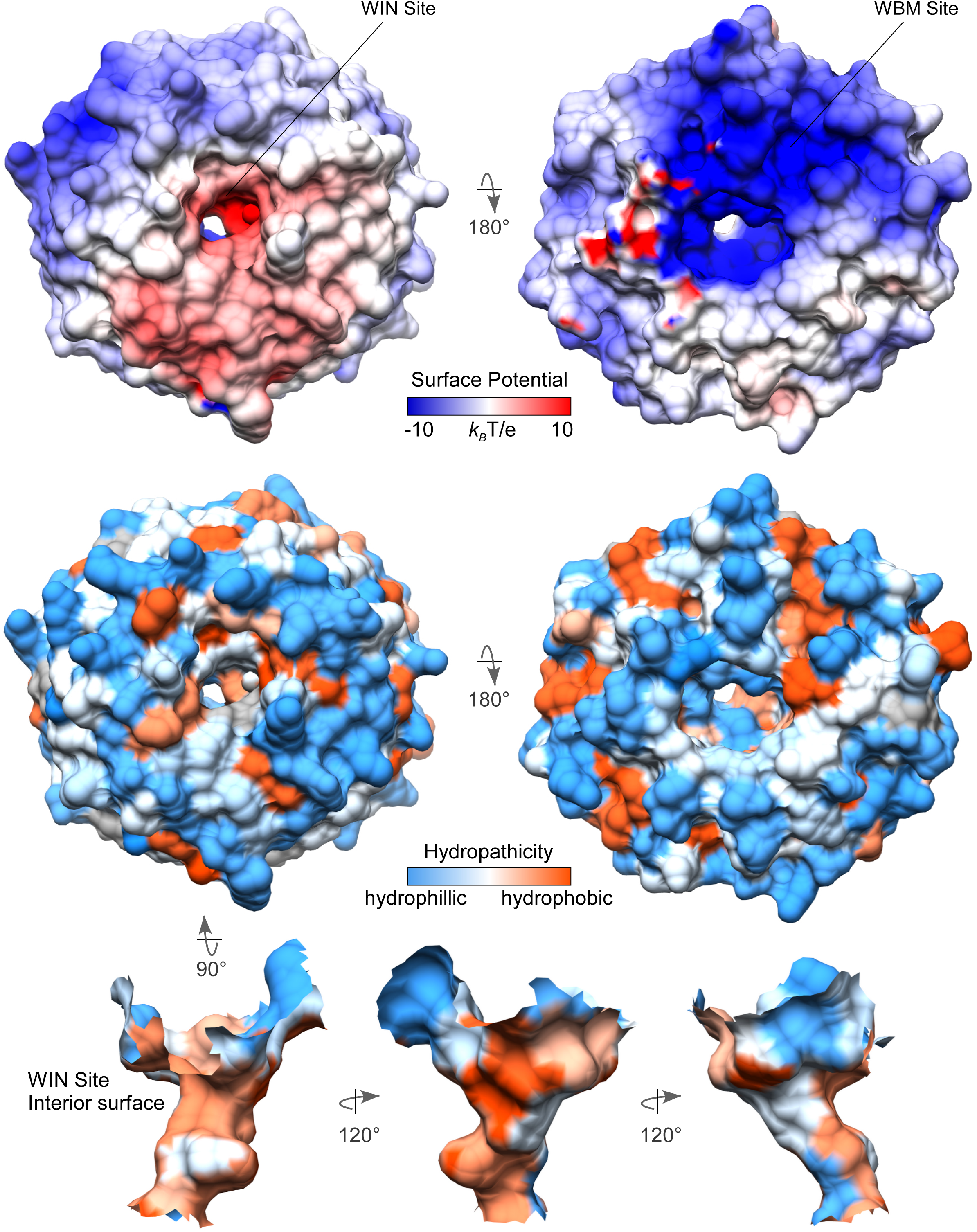
Electrostatic potential (top) and hydropathicity (bottom) surface representations of the WIN site

**Figure S5.**
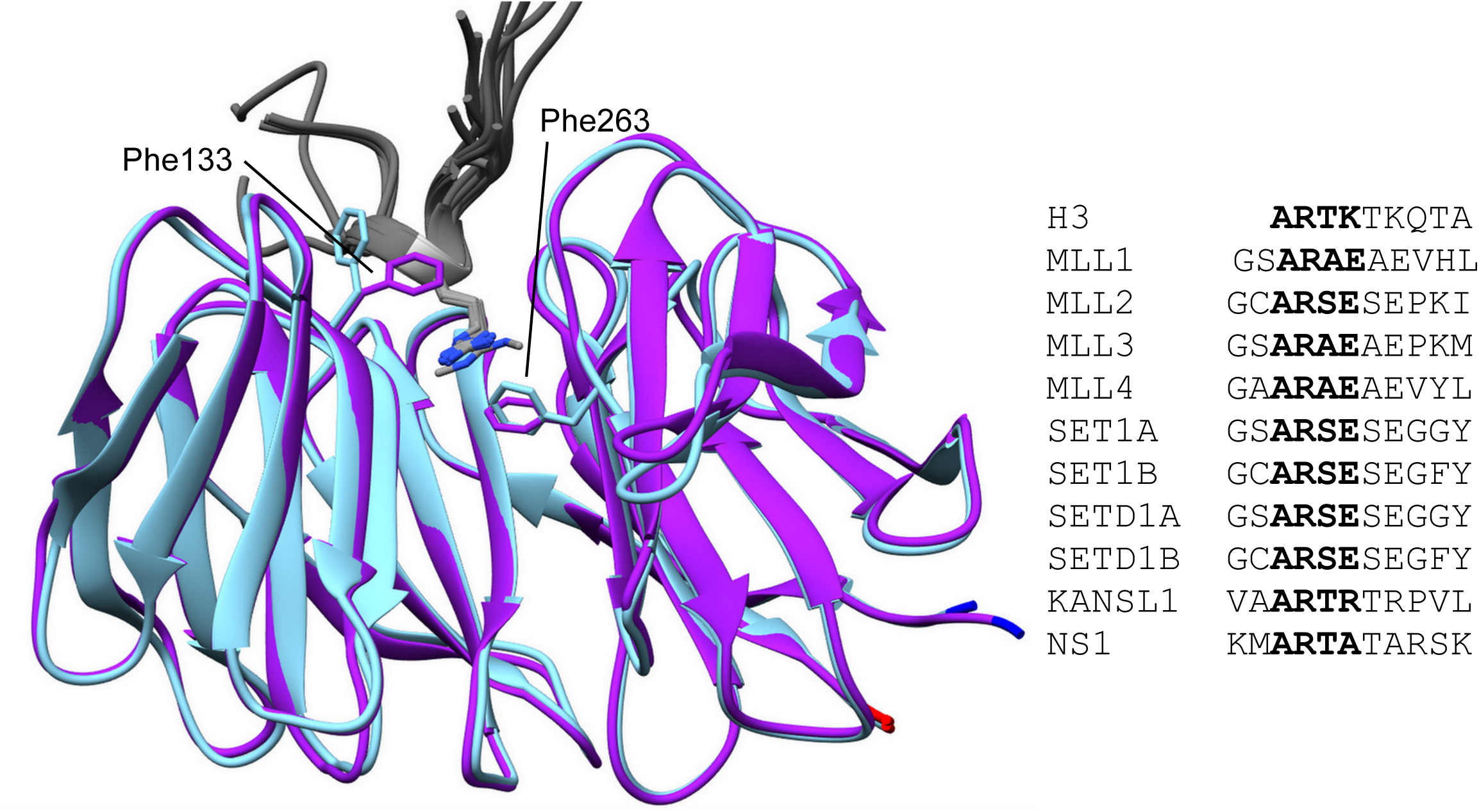
Superposition of apo-WDR5 (cyan) and WDR5 in complex (purple) with structurally characterized WIN site ligands (grey) showing common binding mode with the arginine sidechain guanidino group stacking between WDR5 residues Phe133 and Phe263. Ligand WIN motifs (bolded) are aligned.

**Figure S6.**
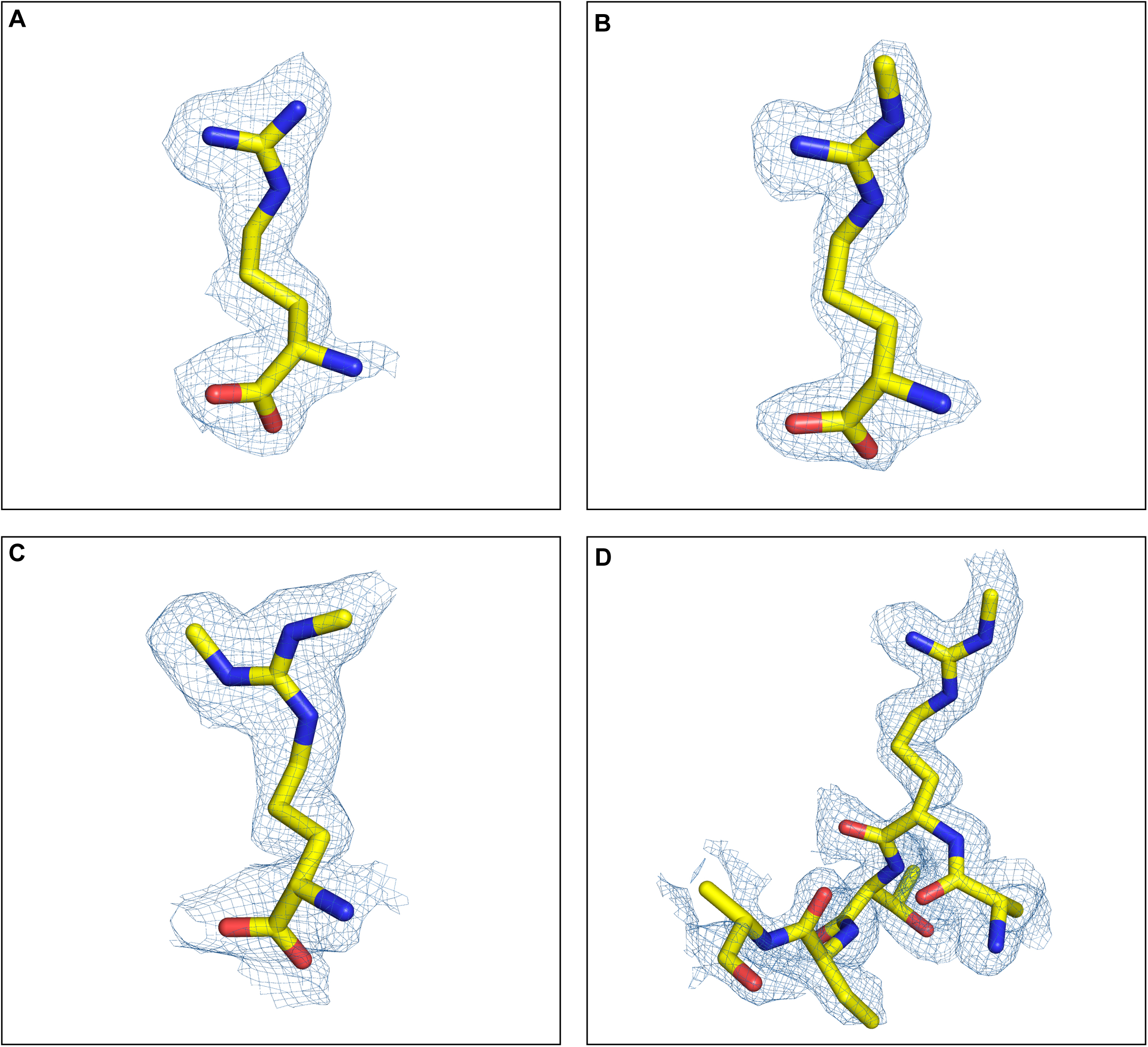
Ligand density maps of **a.** L-Arg **b.** me1-L-Arg **c.** me2s-L-Arg and **d.** H3R2me1 peptide in complex with WDR5

**Figure S7.**
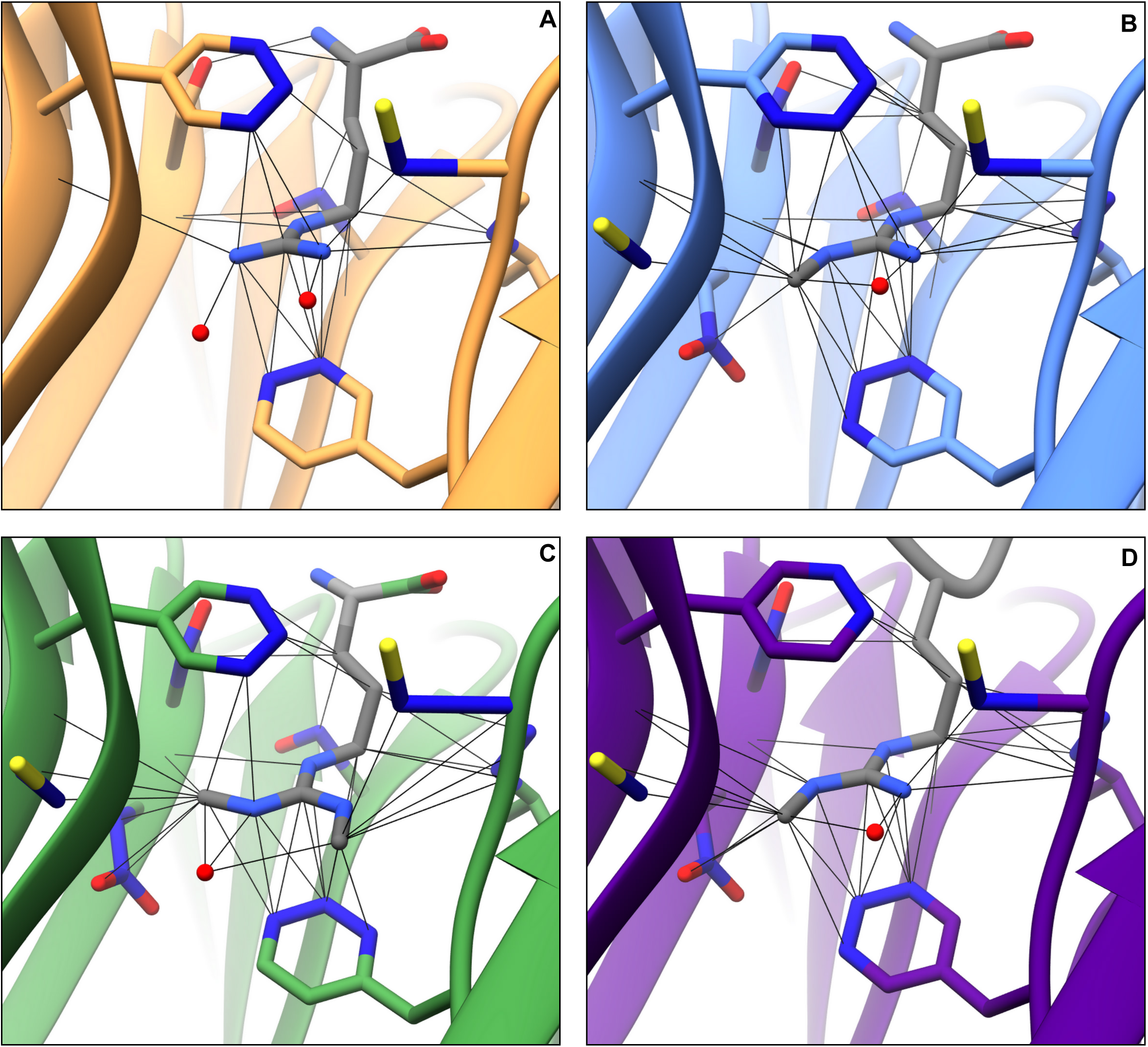
Contacts within VDW distances depicted for **a.** L-Arg, **b.** L-Arg me1, **c.** L-Arg me2s, and **d.** H3R2me1 peptide ligands. Backbone contacts uncolored; contacts with WDR5 residue sidechain atoms are colored dark blue. VDW contacts between L-Arg me2s ω_me_’ and S218 and between L-Arg me2s ω_me_’ and F219 sidechain omitted for clarity. Difference in contacts summarized in Tables 2, S3. H2O (red sphere).

**Figure S8.**
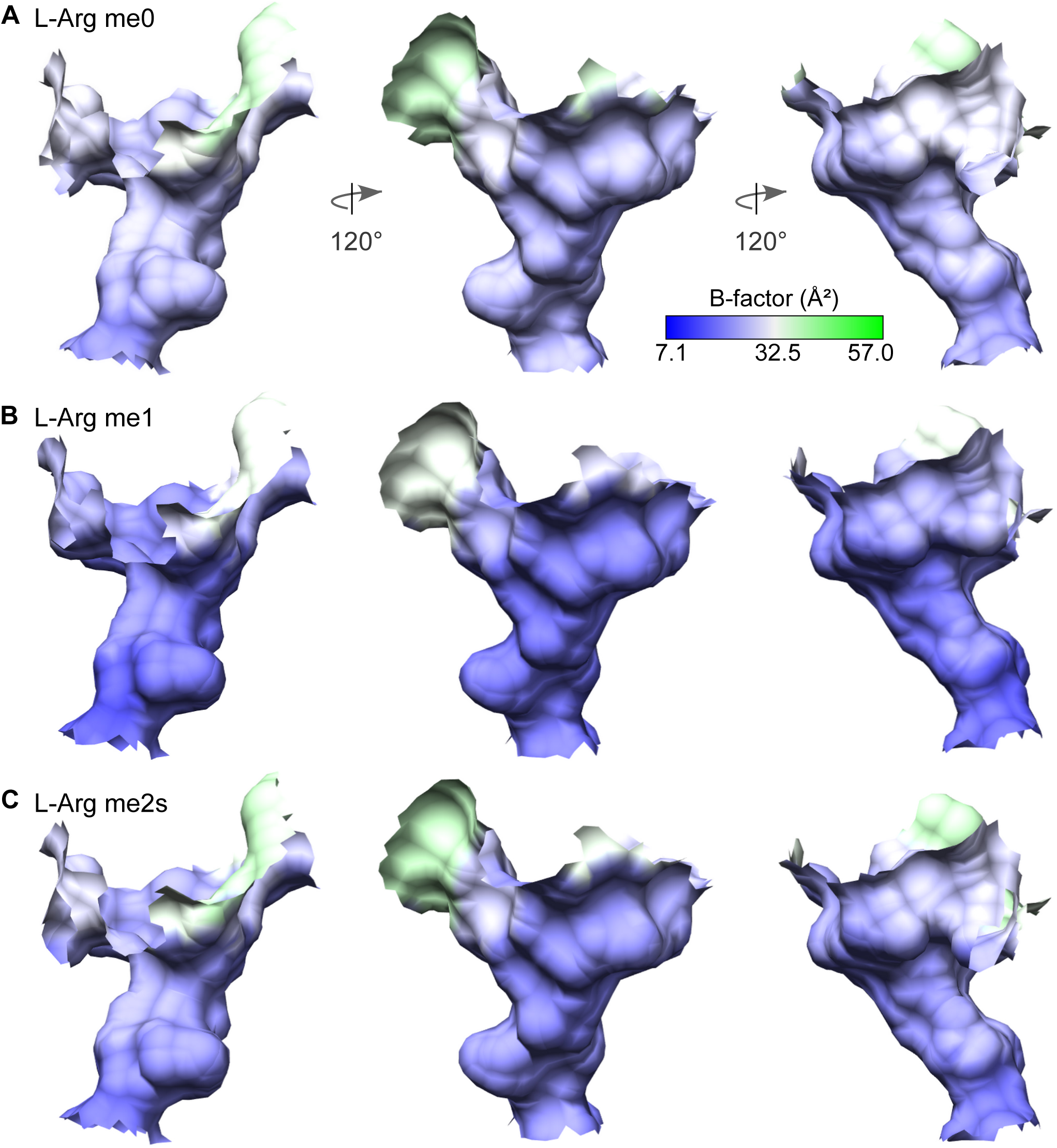
Surface representation of WIN site colored by B-factor showing no major differences when liganded to **a.** L-Arg, **b.** L-Arg me1, and **c.** L-Arg me2s.

**Figure S9.**
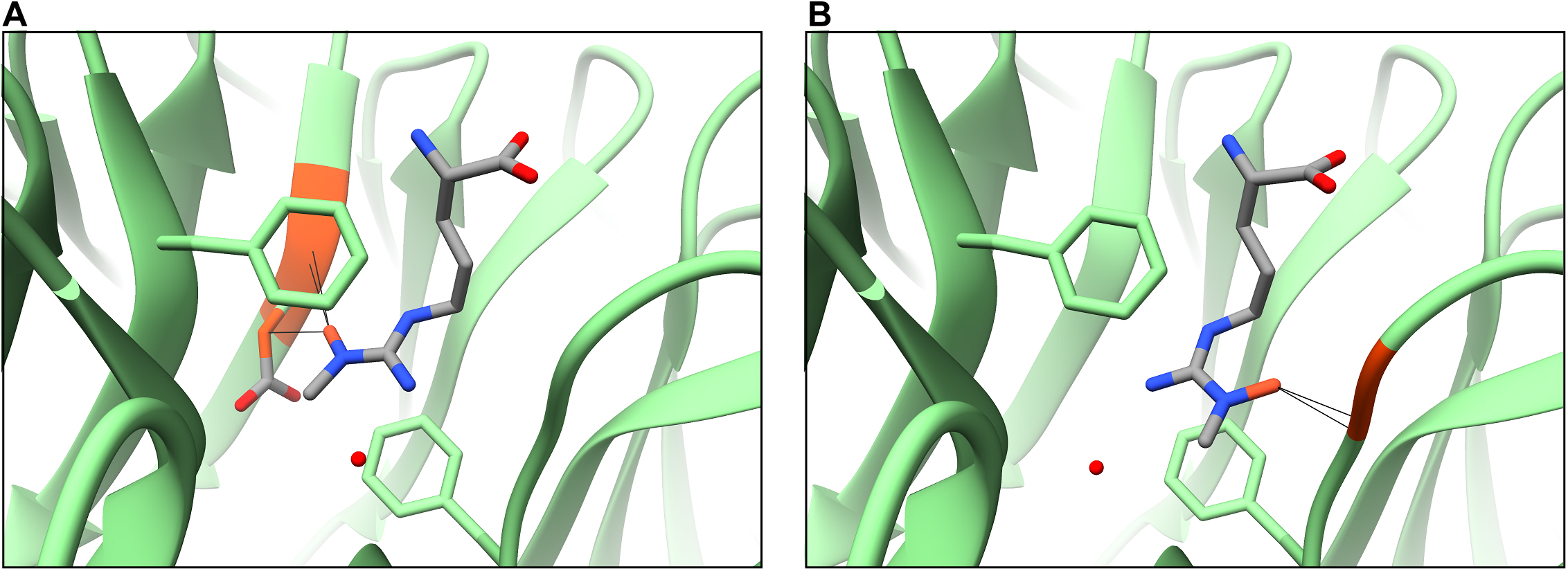
Rme2a modeled in the WIN site depicting clashes that potentially inhibit binding. **a.** Rme2a modeled from Rme1 structure. Clashes with Ser91 backbone and Asp92 sidechain higlighed in orange. **b.** Rme2a modeled from Rme2s structure. Clashes with Cys261 backbone highlighted in orange.

## REFERENCES

1. Gayatri, S.; Bedford, M. T., Readers of histone methylarginine marks. Biochimica et biophysica acta 2014, 1839 (8), 702–10.

2. Morales, Y.; Caceres, T.; May, K.; Hevel, J. M., Biochemistry and regulation of the protein arginine methyltransferases (PRMTs). Archives of biochemistry and biophysics 2016, 590, 138–152.

3. Haghandish, N.; Baldwin, R. M.; Morettin, A.; Dawit, H. T.; Adhikary, H.; Masson, J. Y.; Mazroui, R.; Trinkle-Mulcahy, L.; Cote, J., PRMT7 methylates eukaryotic translation initiation factor 2alpha and regulates its role in stress granule formation. Molecular biology of the cell 2019, 30 (6), 778–793.

4. Fuhrmann, J.; Clancy, K. W.; Thompson, P. R., Chemical biology of protein arginine modifications in epigenetic regulation. Chemical reviews 2015, 115 (11), 5413–61.

5. Walport, L. J.; Hopkinson, R. J.; Chowdhury, R.; Schiller, R.; Ge, W.; Kawamura, A.; Schofield, C. J., Arginine demethylation is catalysed by a subset of JmjC histone lysine demethylases. Nature communications 2016, 7, 11974.

6. Lorton, B. M.; Shechter, D., Cellular consequences of arginine methylation. Cellular and molecular life sciences : CMLS 2019, 76 (15), 2933–2956.

7. Guarnaccia, A. D.; Tansey, W. P., Moonlighting with WDR5: A Cellular Multitasker. Journal of clinical medicine 2018, 7 (2).

8. Iberg, A. N.; Espejo, A.; Cheng, D.; Kim, D.; Michaud-Levesque, J.; Richard, S.; Bedford, M. T., Arginine methylation of the histone H3 tail impedes effector binding. The Journal of biological chemistry 2008, 283 (6), 3006–10.

9. Chen, H.; Lorton, B.; Gupta, V.; Shechter, D., A TGFbeta-PRMT5-MEP50 axis regulates cancer cell invasion through histone H3 and H4 arginine methylation coupled transcriptional activation and repression. Oncogene 2017, 36 (3), 373–386.

10. Kim, D. H.; Tang, Z.; Shimada, M.; Fierz, B.; Houck-Loomis, B.; Bar-Dagen, M.; Lee, S.; Lee, S. K.; Muir, T. W.; Roeder, R. G.; Lee, J. W., Histone H3K27 trimethylation inhibits H3 binding and function of SET1-like H3K4 methyltransferase complexes. Molecular and cellular biology 2013, 33 (24), 4936–46.

11. Schuetz, A.; Allali-Hassani, A.; Martin, F.; Loppnau, P.; Vedadi, M.; Bochkarev, A.; Plotnikov, A. N.; Arrowsmith, C. H.; Min, J., Structural basis for molecular recognition and presentation of histone H3 by WDR5. The EMBO journal 2006, 25 (18), 4245–52.

12. Wysocka, J.; Swigut, T.; Milne, T. A.; Dou, Y.; Zhang, X.; Burlingame, A. L.; Roeder, R. G.; Brivanlou, A. H.; Allis, C. D., WDR5 associates with histone H3 methylated at K4 and is essential for H3 K4 methylation and vertebrate development. Cell 2005, 121 (6), 859–72.

13. Dong, F.; Li, Q.; Yang, C.; Huo, D.; Wang, X.; Ai, C.; Kong, Y.; Sun, X.; Wang, W.; Zhou, Y.; Liu, X.; Li, W.; Gao, W.; Liu, W.; Kang, C.; Wu, X., PRMT2 links histone H3R8 asymmetric dimethylation to oncogenic activation and tumorigenesis of glioblastoma. Nature communications 2018, 9 (1), 4552.

14. Pal, S.; Vishwanath, S. N.; Erdjument-Bromage, H.; Tempst, P.; Sif, S., Human SWI/SNF-associated PRMT5 methylates histone H3 arginine 8 and negatively regulates expression of ST7 and NM23 tumor suppressor genes. Molecular and cellular biology 2004, 24 (21), 9630–45.

15. Ruthenburg, A. J.; Wang, W.; Graybosch, D. M.; Li, H.; Allis, C. D.; Patel, D. J.; Verdine, G. L., Histone H3 recognition and presentation by the WDR5 module of the MLL1 complex. Nature structural & molecular biology 2006, 13 (8), 704–12.

16. Couture, J. F.; Collazo, E.; Trievel, R. C., Molecular recognition of histone H3 by the WD40 protein WDR5. Nature structural & molecular biology 2006, 13 (8), 698–703.

17. Avdic, V.; Zhang, P.; Lanouette, S.; Voronova, A.; Skerjanc, I.; Couture, J. F., Fine-tuning the stimulation of MLL1 methyltransferase activity by a histone H3-based peptide mimetic. FASEB journal : official publication of the Federation of American Societies for Experimental Biology 2011, 25 (3), 960–7.

18. Migliori, V.; Muller, J.; Phalke, S.; Low, D.; Bezzi, M.; Mok, W. C.; Sahu, S. K.; Gunaratne, J.; Capasso, P.; Bassi, C.; Cecatiello, V.; De Marco, A.; Blackstock, W.; Kuznetsov, V.; Amati, B.; Mapelli, M.; Guccione, E., Symmetric dimethylation of H3R2 is a newly identified histone mark that supports euchromatin maintenance. Nature structural & molecular biology 2012, 19 (2), 136–44.

19. Guccione, E.; Bassi, C.; Casadio, F.; Martinato, F.; Cesaroni, M.; Schuchlautz, H.; Luscher, B.; Amati, B., Methylation of histone H3R2 by PRMT6 and H3K4 by an MLL complex are mutually exclusive. Nature 2007, 449 (7164), 933–7.

20. Hyllus, D.; Stein, C.; Schnabel, K.; Schiltz, E.; Imhof, A.; Dou, Y.; Hsieh, J.; Bauer, U. M., PRMT6-mediated methylation of R2 in histone H3 antagonizes H3 K4 trimethylation. Genes & development 2007, 21 (24), 3369–80.

21. Lakowski, T. M.; Frankel, A., A kinetic study of human protein arginine N-methyltransferase 6 reveals a distributive mechanism. The Journal of biological chemistry 2008, 283 (15), 10015–25.

22. Kolbel, K.; Ihling, C.; Bellmann-Sickert, K.; Neundorf, I.; Beck-Sickinger, A. G.; Sinz, A.; Kuhn, U.; Wahle, E., Type I Arginine Methyltransferases PRMT1 and PRMT-3 Act Distributively. The Journal of biological chemistry 2009, 284 (13), 8274–82.

23. Guo, A.; Gu, H.; Zhou, J.; Mulhern, D.; Wang, Y.; Lee, K. A.; Yang, V.; Aguiar, M.; Kornhauser, J.; Jia, X.; Ren, J.; Beausoleil, S. A.; Silva, J. C.; Vemulapalli, V.; Bedford, M. T.; Comb, M. J., Immunoaffinity enrichment and mass spectrometry analysis of protein methylation. Molecular & cellular proteomics : MCP 2014, 13 (1), 372–87.

24. Geoghegan, V.; Guo, A.; Trudgian, D.; Thomas, B.; Acuto, O., Comprehensive identification of arginine methylation in primary T cells reveals regulatory roles in cell signalling. Nature communications 2015, 6, 6758.

25. Larsen, S. C.; Sylvestersen, K. B.; Mund, A.; Lyon, D.; Mullari, M.; Madsen, M. V.; Daniel, J. A.; Jensen, L. J.; Nielsen, M. L., Proteome-wide analysis of arginine monomethylation reveals widespread occurrence in human cells. Science signaling 2016, 9 (443), rs9.

26. Yakubu, R. R.; Silmon de Monerri, N. C.; Nieves, E.; Kim, K.; Weiss, L. M., Comparative Monomethylarginine Proteomics Suggests that Protein Arginine Methyltransferase 1 (PRMT1) is a Significant Contributor to Arginine Monomethylation in Toxoplasma gondii. Molecular & cellular proteomics : MCP 2017, 16 (4), 567–580.

27. Duncan, K. W.; Rioux, N.; Boriack-Sjodin, P. A.; Munchhof, M. J.; Reiter, L. A.; Majer, C. R.; Jin, L.; Johnston, L. D.; Chan-Penebre, E.; Kuplast, K. G.; Porter Scott, M.; Pollock, R. M.; Waters, N. J.; Smith, J. J.; Moyer, M. P.; Copeland, R. A.; Chesworth, R., Structure and Property Guided Design in the Identification of PRMT5 Tool Compound EPZ015666. ACS medicinal chemistry letters 2016, 7 (2), 162–6.

28. Feng, Y.; Wang, J.; Asher, S.; Hoang, L.; Guardiani, C.; Ivanov, I.; Zheng, Y. G., Histone H4 acetylation differentially modulates arginine methylation by an in Cis mechanism. The Journal of biological chemistry 2011, 286 (23), 20323–34.

29. Battye, T. G.; Kontogiannis, L.; Johnson, O.; Powell, H. R.; Leslie, A. G., iMOSFLM: a new graphical interface for diffraction-image processing with MOSFLM. Acta crystallographica. Section D, Biological crystallography 2011, 67 (Pt 4), 271–81.

30. The CCP4 suite: programs for protein crystallography. Acta crystallographica. Section D, Biological crystallography 1994, 50 (Pt 5), 760–3.

31. Adams, P. D.; Afonine, P. V.; Bunkoczi, G.; Chen, V. B.; Davis, I. W.; Echols, N.; Headd, J. J.; Hung, L. W.; Kapral, G. J.; Grosse-Kunstleve, R. W.; McCoy, A. J.; Moriarty, N. W.; Oeffner, R.; Read, R. J.; Richardson, D. C.; Richardson, J. S.; Terwilliger, T. C.; Zwart, P. H., PHENIX: a comprehensive Python-based system for macromolecular structure solution. *Acta crystallographica. Section D*, Biological crystallography 2010, 66 (Pt 2), 213–21.

32. McCoy, A. J.; Grosse-Kunstleve, R. W.; Adams, P. D.; Winn, M. D.; Storoni, L. C.; Read, R. J., Phaser crystallographic software. Journal of applied crystallography 2007, 40 (Pt 4), 658–674.

33. Emsley, P.; Lohkamp, B.; Scott, W. G.; Cowtan, K., Features and development of Coot. Acta crystallographica. Section D, Biological crystallography 2010, 66 (Pt 4), 486–501.

34. Murshudov, G. N.; Vagin, A. A.; Dodson, E. J., Refinement of macromolecular structures by the maximum-likelihood method. Acta crystallographica. Section D, Biological crystallography 1997, 53 (Pt 3), 240–55.

35. Pettersen, E. F.; Goddard, T. D.; Huang, C. C.; Couch, G. S.; Greenblatt, D. M.; Meng, E. C.; Ferrin, T. E., UCSF Chimera--a visualization system for exploratory research and analysis. Journal of computational chemistry 2004, 25 (13), 1605–12.

36. Chen, V. B.; Arendall, W. B., 3rd; Headd, J. J.; Keedy, D. A.; Immormino, R. M.; Kapral, G. J.; Murray, L. W.; Richardson, J. S.; Richardson, D. C., MolProbity: all-atom structure validation for macromolecular crystallography. *Acta crystallographica. Section D*, Biological crystallography 2010, 66 (Pt 1), 12–21.

37. Goldfarb, A. R.; Saidel, L. J.; Mosovich, E., The ultraviolet absorption spectra of proteins. The Journal of biological chemistry 1951, 193 (1), 397–404.

38. Anthis, N. J.; Clore, G. M., Sequence-specific determination of protein and peptide concentrations by absorbance at 205 nm. Protein science : a publication of the Protein Society 2013, 22 (6), 851–8.

39. Scopes, R. K., Measurement of protein by spectrophotometry at 205 nm. Analytical biochemistry 1974, 59 (1), 277–82.

40. Grebien, F.; Vedadi, M.; Getlik, M.; Giambruno, R.; Grover, A.; Avellino, R.; Skucha, A.; Vittori, S.; Kuznetsova, E.; Smil, D.; Barsyte-Lovejoy, D.; Li, F.; Poda, G.; Schapira, M.; Wu, H.; Dong, A.; Senisterra, G.; Stukalov, A.; Huber, K. V. M.; Schonegger, A.; Marcellus, R.; Bilban, M.; Bock, C.; Brown, P. J.; Zuber, J.; Bennett, K. L.; Al-Awar, R.; Delwel, R.; Nerlov, C.; Arrowsmith, C. H.; Superti-Furga, G., Pharmacological targeting of the Wdr5-MLL interaction in C/EBPalpha N-terminal leukemia. Nature chemical biology 2015, 11 (8), 571–578.

41. Li, D. D.; Wang, Z. H.; Chen, W. L.; Xie, Y. Y.; You, Q. D.; Guo, X. K., Structure-based design of ester compounds to inhibit MLL complex catalytic activity by targeting mixed lineage leukemia 1 (MLL1)-WDR5 interaction. Bioorganic & medicinal chemistry 2016, 24 (22), 6109–6118.

42. Huggins, D. J., Quantifying the entropy of binding for water molecules in protein cavities by computing correlations. Biophysical journal 2015, 108 (4), 928–936.

43. Xue, H.; Yao, T.; Cao, M.; Zhu, G.; Li, Y.; Yuan, G.; Chen, Y.; Lei, M.; Huang, J., Structural basis of nucleosome recognition and modification by MLL methyltransferases. Nature 2019, 573 (7774), 445–449.

44. Park, S. H.; Ayoub, A.; Lee, Y. T.; Xu, J.; Kim, H.; Zheng, W.; Zhang, B.; Sha, L.; An, S.; Zhang, Y.; Cianfrocco, M. A.; Su, M.; Dou, Y.; Cho, U. S., Cryo-EM structure of the human MLL1 core complex bound to the nucleosome. Nature communications 2019, 10 (1), 5540.

45. Patel, A.; Dharmarajan, V.; Cosgrove, M. S., Structure of WDR5 bound to mixed lineage leukemia protein-1 peptide. The Journal of biological chemistry 2008, 283 (47), 32158–61.

46. Song, J. J.; Kingston, R. E., WDR5 interacts with mixed lineage leukemia (MLL) protein via the histone H3-binding pocket. The Journal of biological chemistry 2008, 283 (50), 35258–64.

47. Dharmarajan, V.; Lee, J. H.; Patel, A.; Skalnik, D. G.; Cosgrove, M. S., Structural basis for WDR5 interaction (Win) motif recognition in human SET1 family histone methyltransferases. The Journal of biological chemistry 2012, 287 (33), 27275–89.

48. Zhang, P.; Lee, H.; Brunzelle, J. S.; Couture, J. F., The plasticity of WDR5 peptide-binding cleft enables the binding of the SET1 family of histone methyltransferases. Nucleic acids research 2012, 40 (9), 4237–46.

49. Dias, J.; Van Nguyen, N.; Georgiev, P.; Gaub, A.; Brettschneider, J.; Cusack, S.; Kadlec, J.; Akhtar, A., Structural analysis of the KANSL1/WDR5/KANSL2 complex reveals that WDR5 is required for efficient assembly and chromatin targeting of the NSL complex. Genes & development 2014, 28 (9), 929–42.

50. Qin, S.; Liu, Y.; Tempel, W.; Eram, M. S.; Bian, C.; Liu, K.; Senisterra, G.; Crombet, L.; Vedadi, M.; Min, J., Structural basis for histone mi *tions* 20micry and hijacking of host proteins by influenza virus protein NS1. Nature communica14, 5, 3952.

