## Supplemental Figures for "A binary arginine methylation switch on histone H3 Arginine 2 regulates its interaction with WDR5"

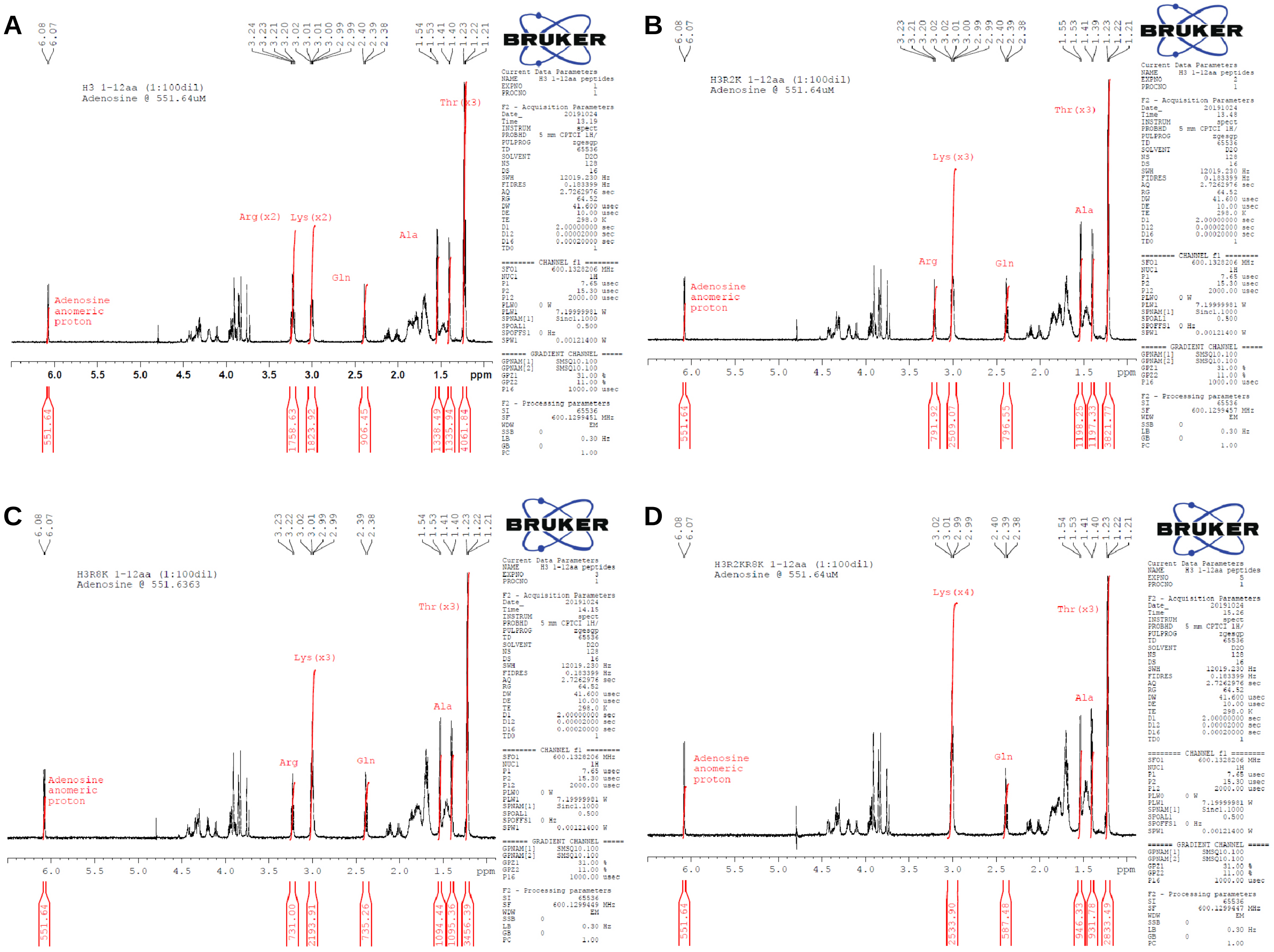


**Figure S1**. 1D $^{1}$H NMR spectra of H3 1-12aa peptides showing peak integrations of residues used to measure peptide concentration. **a.** H3 unmodified **b.** H3R2K **c.** H3R8K. **d.** H3R2KR8K

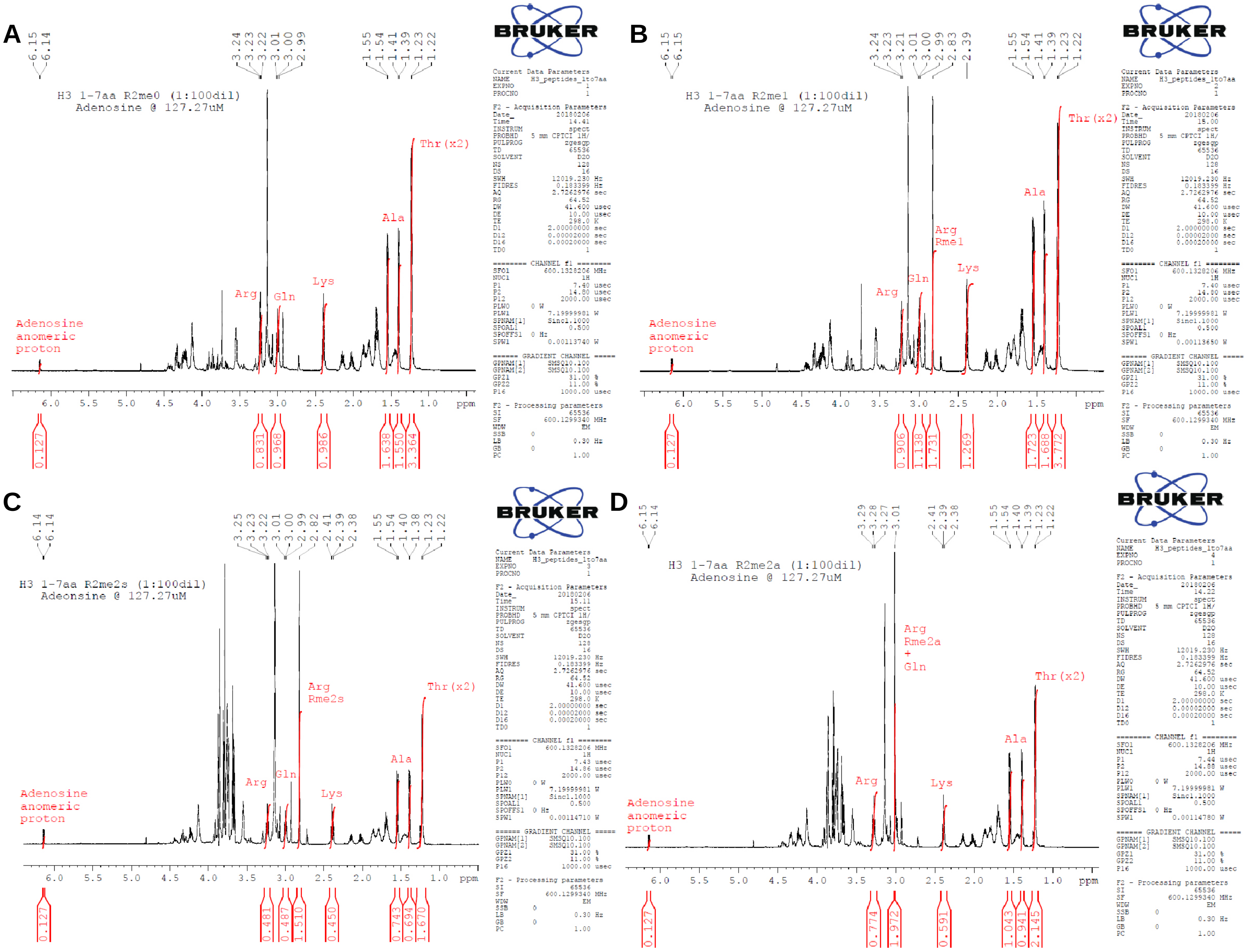


**Figure S2**. 1D $^{1}$H NMR spectra of H3 1-7aa peptides showing peak integrations of residues used to measure peptide concentration. **a.** H3 unmodified **b.** H3R2me1 **c.** H3R2me2s. **d.** H3R2me2a

**A** H3: $NH_2$-ARTKQTARKSTGGKAPRKQLA(GGK-biotin)

Histone H3R2me (1-21aa)
biotinylated peptide pulldown

**B**

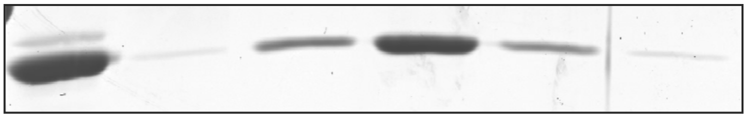


**C**

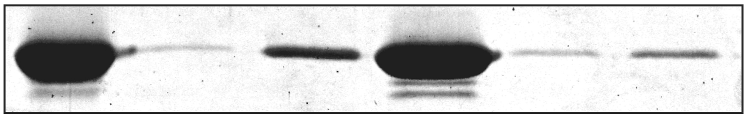


**Figure S3**. Replicate peptide pulldown assays showing WDR5 interacts with H3R2me0, me1, and me2s but not H3R2me2a. (-) negative control: no peptide, resin only. **a.** Histone H3 1-21aa peptide sequence with methylarginine isoforms ocurring at Arg2, underlined. **b.** and **c.** Replicate pulldowns 1 and 2

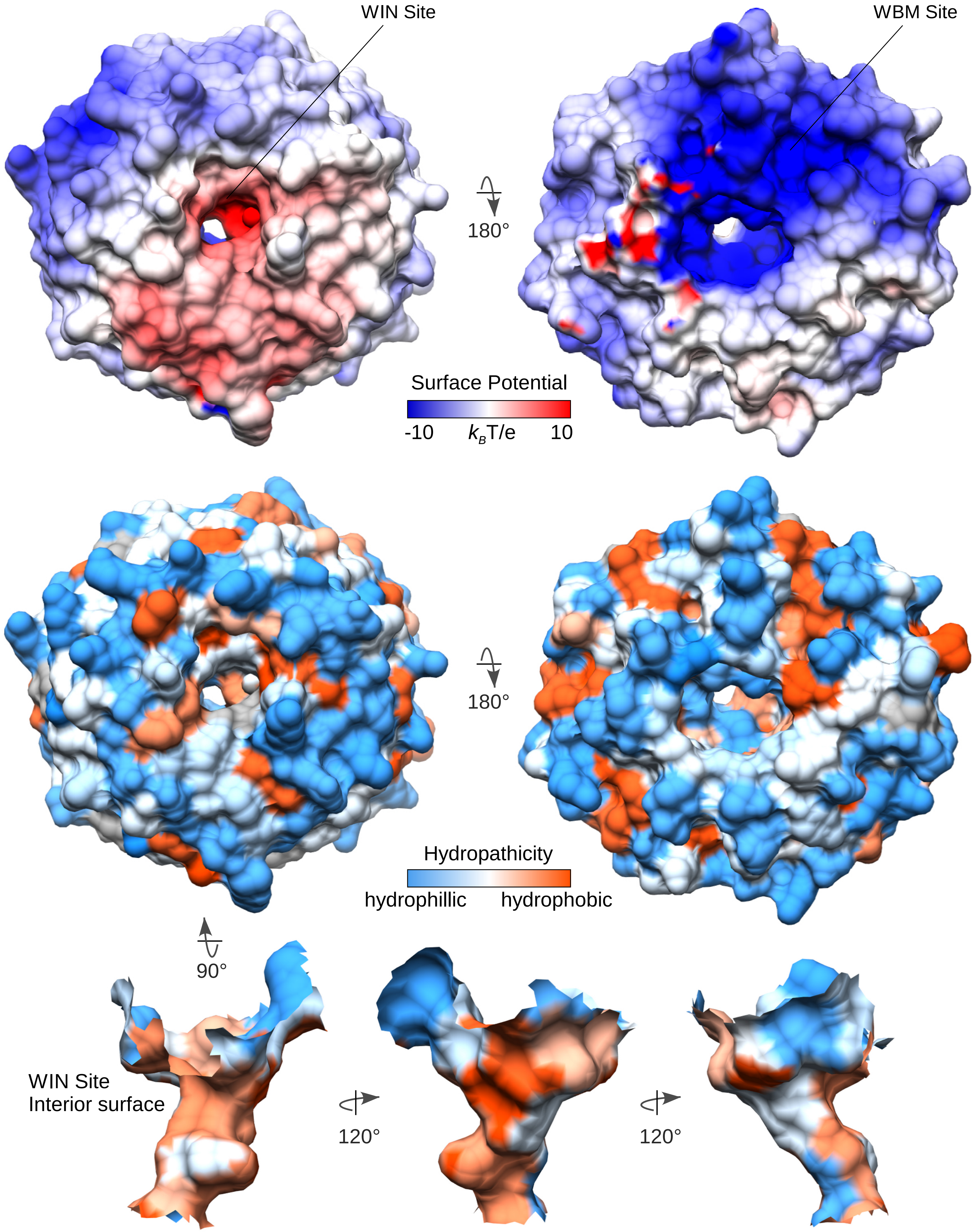


**Figure S4.** Electrostatic potential (top) and hydropathicity (bottom) surface representations of the WIN site

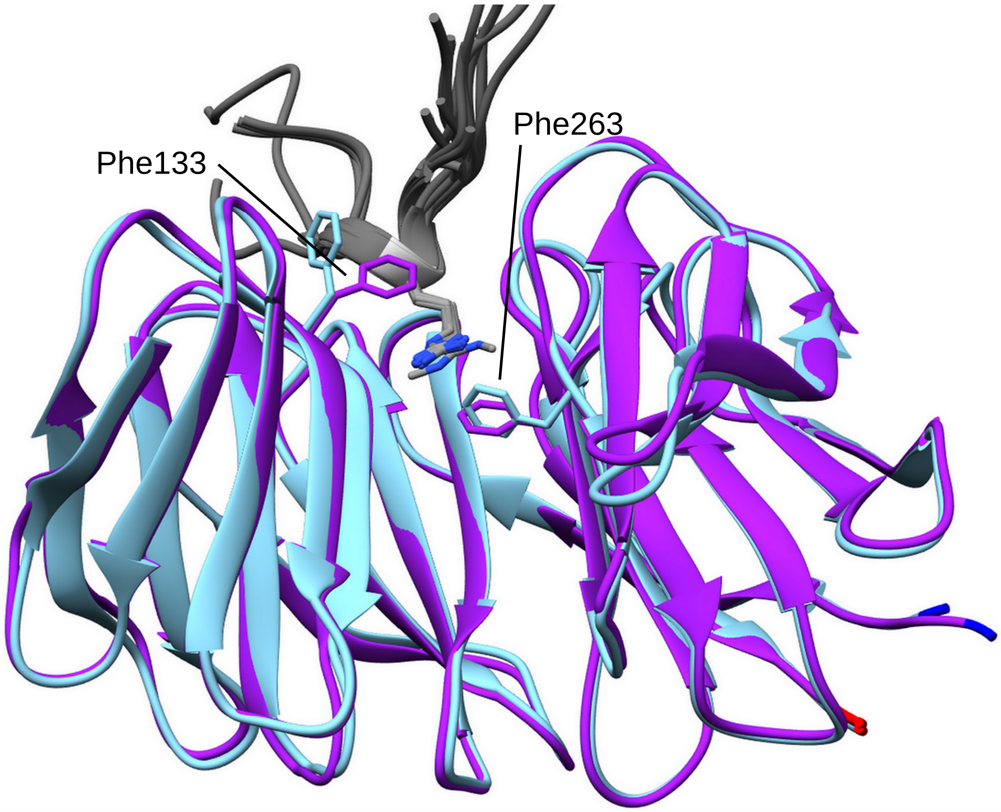


| | |
|---|---|
| H3 | **ARTK**TKQTA |
| MLL1 | GS**ARAE**AEVHL |
| MLL2 | GC**ARSE**SEPKI |
| MLL3 | GS**ARAE**AEPKM |
| MLL4 | GA**ARAE**AEVYL |
| SET1A | GS**ARSE**SEGGY |
| SET1B | GC**ARSE**SEGFY |
| SETD1A | GS**ARSE**SEGGY |
| SETD1B | GC**ARSE**SEGFY |
| KANSL1 | VA**ARTR**TRPVL |
| NS1 | KM**ARTA**TARSK |

**Figure S5.** Superposition of apo-WDR5 (cyan) and WDR5 in complex (purple) with structurally characterized WIN site ligands (grey) showing common binding mode with the arginine sidechain guanidino group stacking between WDR5 residues Phe133 and Phe263. Ligand WIN motifs (bolded) are aligned.

**Figure S6:** Ligand electron density

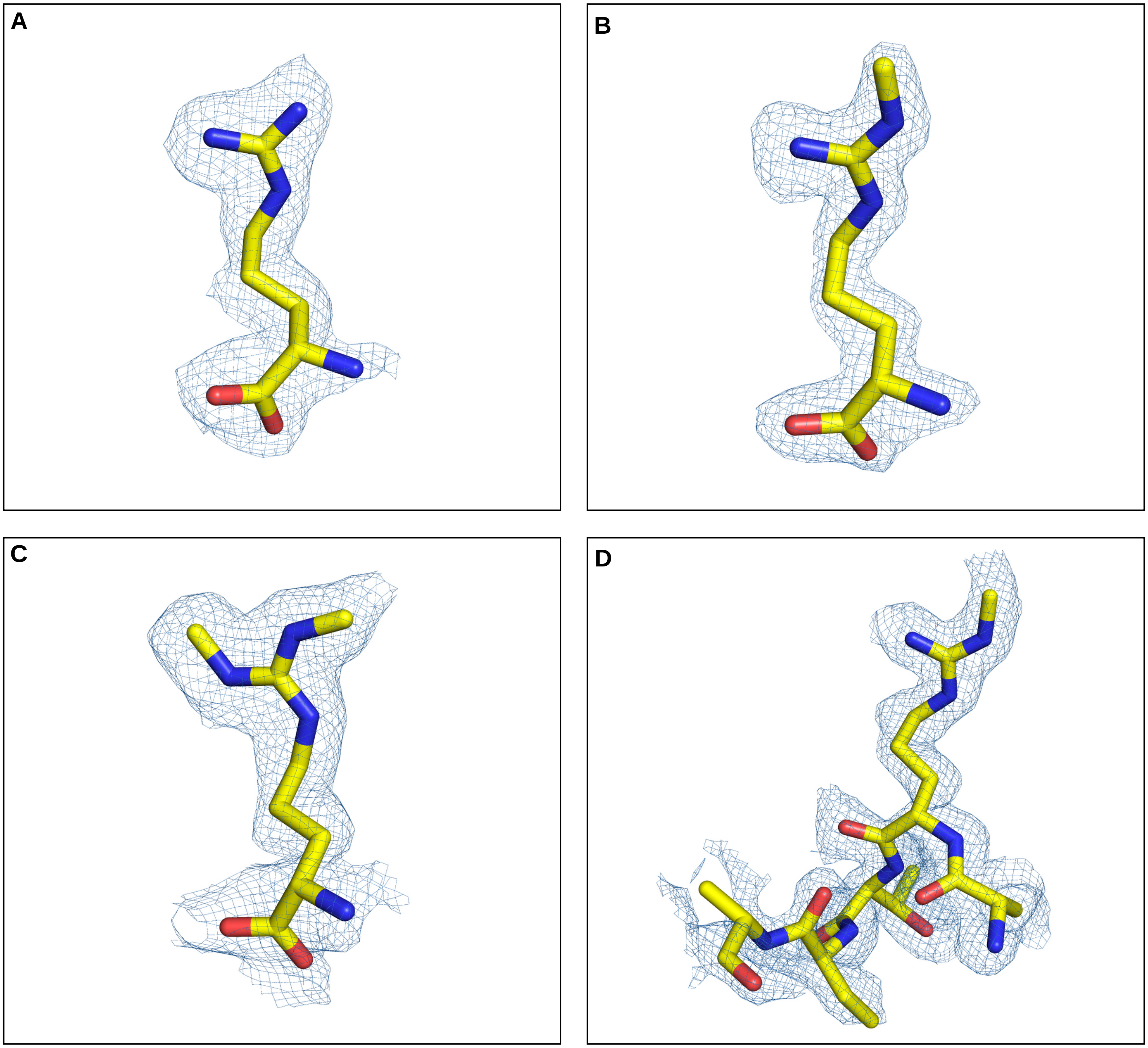


**Figure S6.** Ligand density maps of **a.** L-Arg **b.** me1-L-Arg **c.** me2s-L-Arg and **d.** H3R2me1 peptide in complex with WDR5

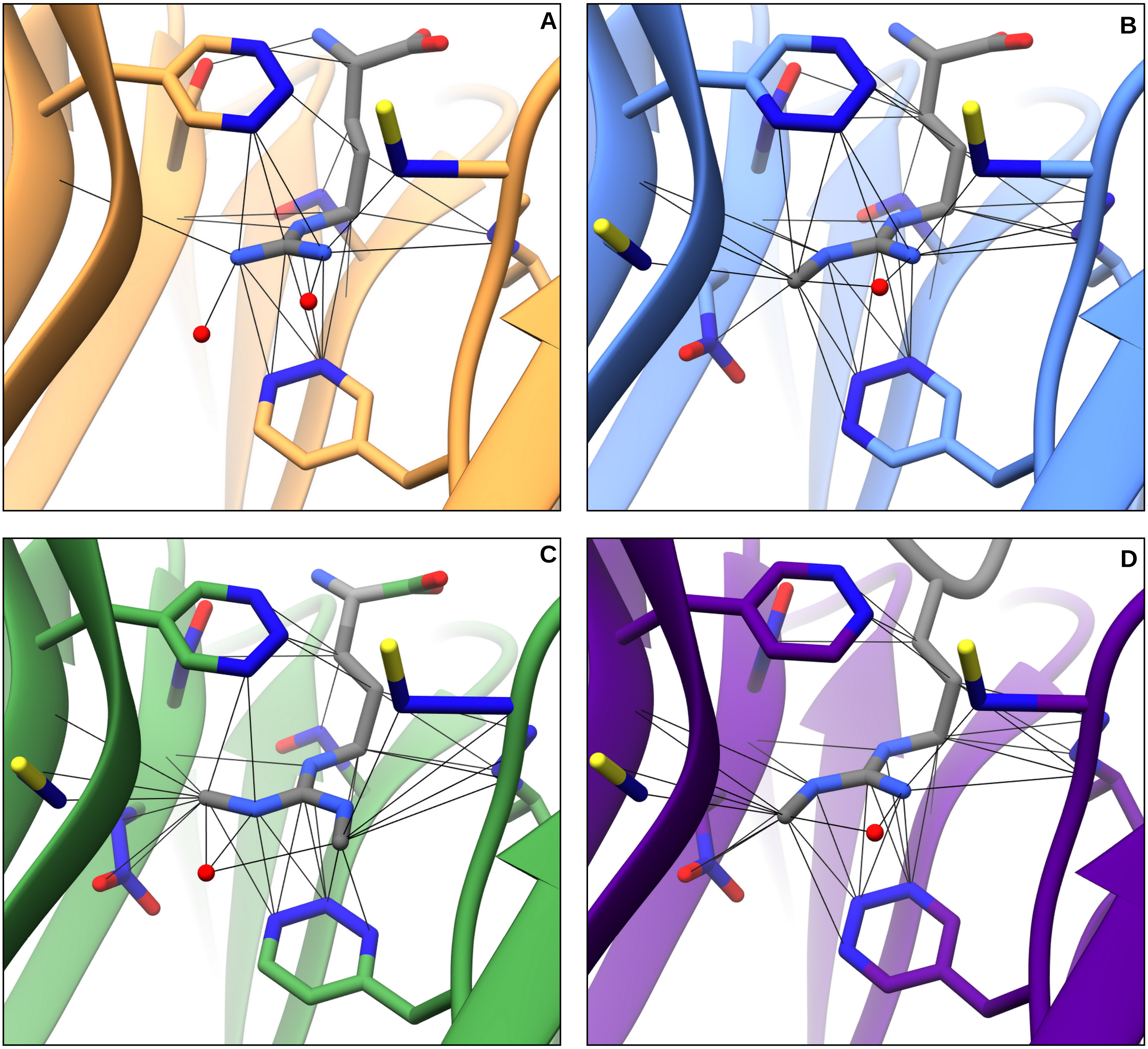


**Figure S7.** Contacts within VDW distances depicted for **a.** L-Arg, **b.** L-Arg me1, **c.** L-Arg me2s, and **d.** H3R2me1 peptide ligands. Backbone contacts uncolored; contacts with WDR5 residue sidechain atoms are colored dark blue. VDW contacts between L-Arg me2s ωme' and S218 and between L-Arg me2s ωme' and F219 sidechain omitted for clarity. Difference in contacts summarized in Tables 2, S3. $H_2O$ (red sphere).

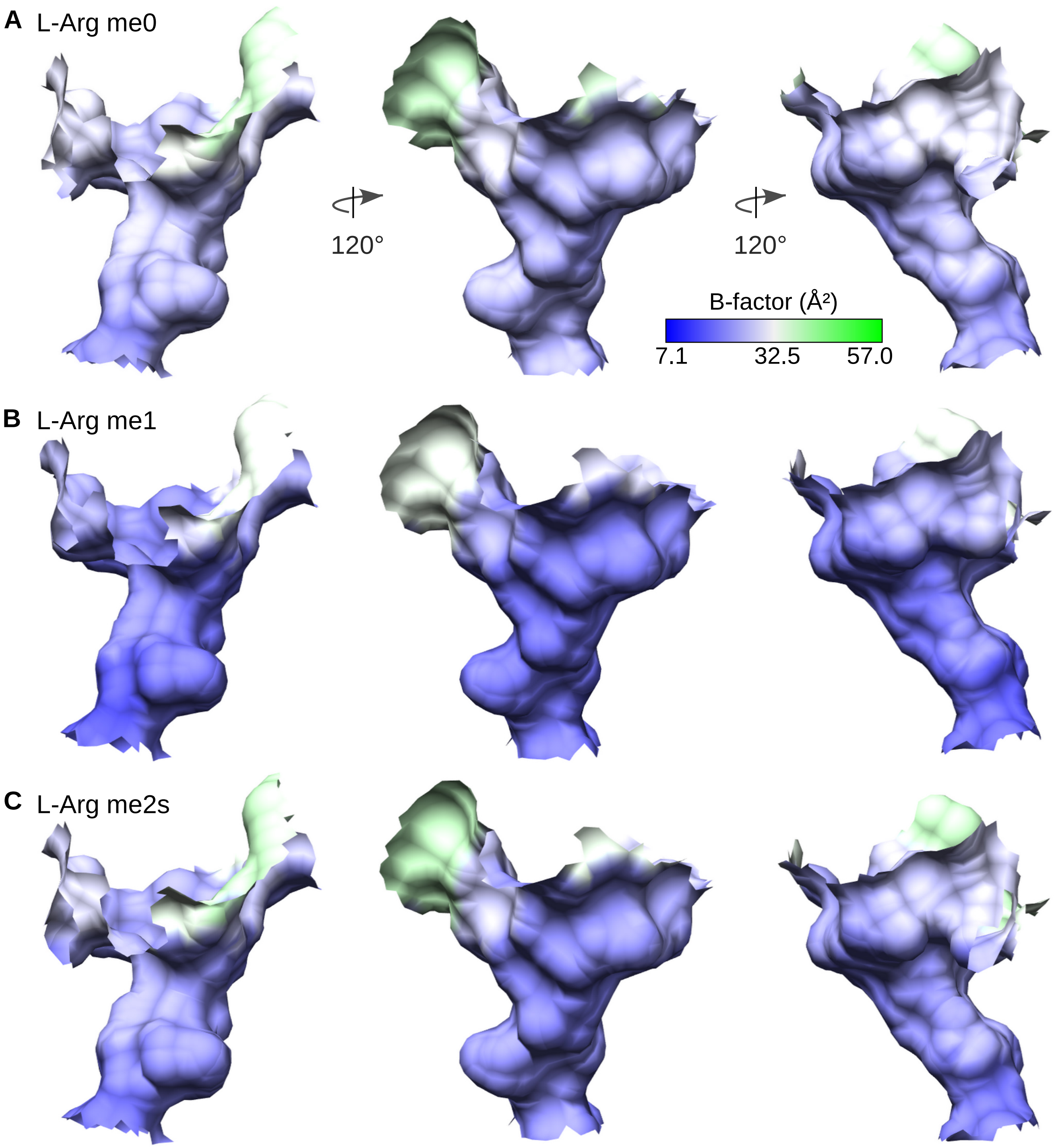


**Figure S8.** Surface representation of WIN site colored by B-factor showing no major differences when liganded to **a.** L-Arg, **b.** L-Arg me1, and **c.** L-Arg me2s.

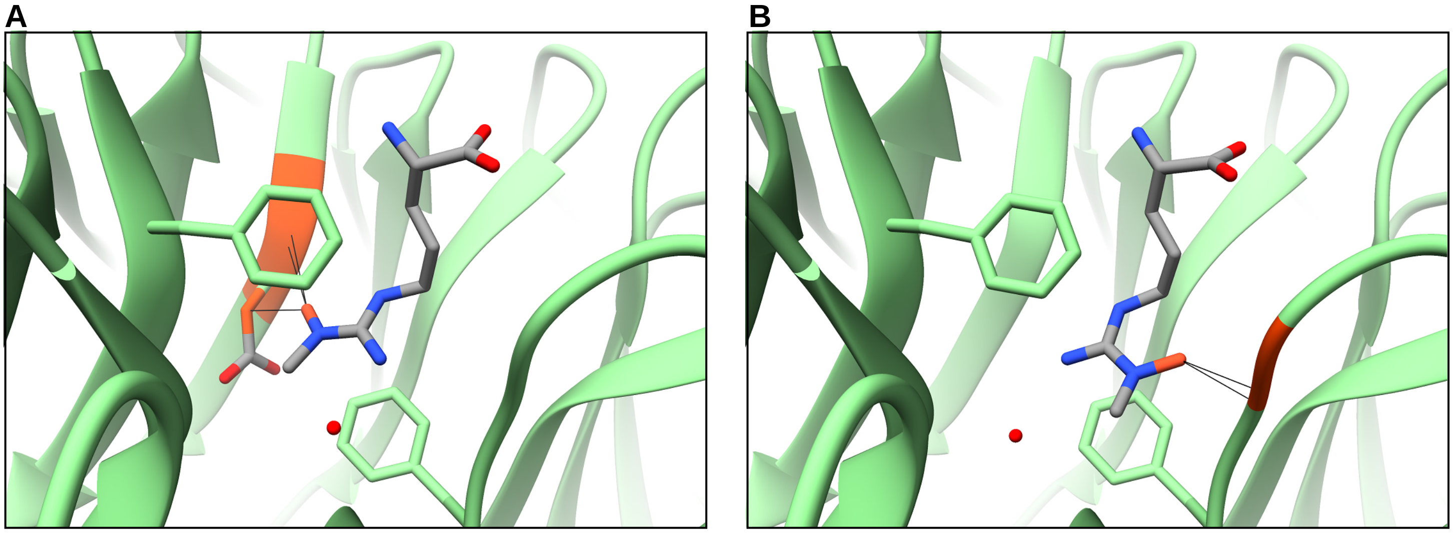


**Figure S9.** Rme2a modeled in the WIN site depicting clashes that potentially inhibit binding. **a.** Rme2a modeled from Rme1 structure. Clashes with Ser91 backbone and Asp92 sidechain higlighed in orange. **b.** Rme2a modeled from Rme2s structure. Clashes with Cys261 backbone highlighted in orange.
